# A genome-wide One Health study of *Klebsiella pneumoniae* in Norway reveals overlapping populations but few recent transmission events across reservoirs

**DOI:** 10.1101/2024.09.11.612360

**Authors:** Marit A K Hetland, Mia A Winkler, Hako Kaspersen, Fredrik Håkonsholm, Ragna-Johanne Bakksjø, Eva Bernhoff, Jose F. Delgado-Blas, Sylvain Brisse, Annapaula Correia, Aasmund Fostervold, Margaret M C Lam, Bjørn-Tore Lunestad, Nachiket P Marathe, Niclas Raffelsberger, Ørjan Samuelsen, Marianne Sunde, Arnfinn Sundsfjord, Anne Margrete Urdahl, Ryan R Wick, Iren H Löhr, Kathryn E Holt

## Abstract

Members of the *Klebsiella pneumoniae* species complex (KpSC) are opportunistic pathogens that cause severe and difficult-to-treat infections. KpSC are common in non-human niches, but the clinical relevance of these populations is disputed. Utilising 3,255 whole-genome sequenced isolates from human, animal and marine sources collected during 2001-2020 in Norway, we showed the KpSC populations in different niches were distinct but overlapping. Notably, human infection isolates showed greatest connectivity with each other, followed by isolates from human carriage, pigs, and bivalves. Nearly 5% of human infection isolates had close relatives (≤22 substitutions) amongst animal and marine isolates, despite temporally and geographically distant sampling of these sources. Infection prevention measures are essential to limit transmission within human clinical settings and reduce disease burden. However, as colonisation often precedes infection, preventing transmission that leads to colonisation, e.g. transmission between animals and humans in the community, and in the food chain, could also be beneficial.

## Introduction

*Klebsiella pneumoniae* are a frequent cause of difficult-to-treat multidrug resistant (MDR) infections. Additionally, strains with acquired virulence factors can cause severe hypervirulent infections, and convergent strains displaying both MDR and hypervirulent traits are rapidly emerging ^1^. The *K. pneumoniae* species complex (KpSC) consists of seven species/subspecies that can be further divided into sublineages (SLs) and sequence types (STs) ^2,3^. The KpSC are opportunistic pathogens that often colonise people before subsequently causing infections ^4^. They can also be found in environmental sources, including terrestrial and marine animals and food, which could provide a source of transmission to humans ^5,6^. However, it has been unclear to what extent KpSC populations isolated from humans are distinct from those found in other niches, and whether non-human sources serve as reservoirs for clinically relevant strains, contributing to the pool of bacteria that cause colonisation and infection in humans. For example, depending on the availability of nutrients and exposure to substances such as antibiotics and heavy metals, KpSC from different niches may display distinct metabolic traits and antimicrobial resistance (AMR) levels due to selective pressures and niche adaptation ^7,8^. By identifying niche-specific traits, we may better understand the ecological distribution of strains, and distinguish which KpSC niches represent clinically relevant reservoirs of infections in humans versus those with little relevance to human health.

To study the emergence, dynamics and spread of KpSC within and between niches, a One Health approach can be applied, using whole-genome sequencing to characterise and compare bacterial isolates from diverse sources. The results from such studies have the potential to guide public health interventions, by identifying routes of transmission that could be interrupted to reduce infections, or by identifying niches where selection for clinically relevant AMR or virulence is occurring.

Only a few One Health studies of KpSC have previously been reported, and few (if any) have assessed niche-adaptation in KpSC. These largely concluded that short-term transmission of KpSC is more common within clinical settings than between clinical and other sources, although transmission to humans from other sources does occasionally occur ^5,9,10^. This is in contrast to classic foodborne bacterial pathogens such as *Salmonella enterica* or *Campylobacter jejuni*, where human infections tend to result from short-chain transmission from animal reservoirs via consumption of contaminated food ^11^. Still, even if KpSC spillover events occur only rarely, they can have significant impacts on human health. To take some well-known examples from virology, the COVID-19 pandemic and MERS epidemic are thought to stem from very rare but high-impact zoonotic transmission from animals to humans, followed by sustained human-to-human transmission ^12,13^. Given that KpSC – unlike *S. enterica* or *C. jejuni* – can cause persistent colonisation of humans and can easily spread between humans, at least in healthcare settings, it is reasonable to propose that even occasional transmission of AMR or hypervirulent KpSC strains from animal reservoirs to humans could have sustained impacts on human health by introducing new clinically relevant strains that subsequently become established in humans.

Since the vast majority of publicly available KpSC genome sequences are derived from humans, there is insufficient data to ascertain whether any of the lineages that are common in humans have an animal origin. However, there are known examples of recent spillover of clinically relevant mobile genetic elements (MGEs) from animals to humans, which have since spread within human-associated KpSC populations. For instance, the *mcr-1* gene, which confers resistance to the last-line antibiotic colistin, is believed to have originated in animals and subsequently spilled over to humans, where it now contributes to difficult-to-treat KpSC and other Enterobacterales infections ^14^. Another example is the pig-associated aerobactin virulence locus *iuc*3, which has been found in multiple KpSC lineages colonising humans and causing infections ^15,16^. It is therefore important to understand how clinically relevant KpSC strains and MGEs enter and circulate among people to inform and support public health intervention measures to prevent disease, health care burdens and mortality.

Investigation of transmission between niches requires sympatric sampling, which has been extremely limited for KpSC ^5,9,17,18^. Through national surveillance programs in the human and veterinary sectors, and screening of human community carriage, animals and marine samples, >3,000 KpSC isolates from across Norway over 20 years have been collected and sequenced ^15,19–23^. The isolates were originally sampled to quantify the burden of infection or carriage in distinct niches, without bias towards AMR or hypervirulence phenotypes. Due to the low prevalence of AMR in Norway ^24^, this provides a unique opportunity to study niche connectivity in a population largely unaffected by antibiotic selection pressures. Here, we report on the diversity and niche-association of strains, genes and other clinically relevant genetic features across a collection of 3,255 KpSC genomes from human, animal and marine niches, and explore evidence for the separation and connectivity of KpSC populations isolated from these niches.

## Results

### KpSC were present among all sources collected on land and in coastal waters

We analysed 3,255 KpSC genomes in this study. Of those, 2,172 (66.7%) were from human infections (n=1,920 blood; 252 urine) from all 22 Norwegian clinical microbiology laboratories between 2001-2018 ^19,23^. Twenty genomes (0.6%) were from animal infections (2018-2020; 13 dog, 4 turkey, 3 broiler). Isolates from 2015-2020 from human and animal carriage studies in Norway were also included (n=1,063, 32.6%). As previously reported, carriage rates estimated in these studies were 16% in humans ^20^ and 0-69% in different animals, with the highest rates in poultry flocks and pigs (Table 1) ^15,21,22^. In addition, KpSC were found in 41.2% (7/17) of surface seawater samples. All isolates were short-read sequenced and 16.9% (n=550) were additionally long-read sequenced. The genomes were divided into three ecological niches (human, animal and marine) and eight source types for comparisons (Fig. 1A).

**Fig. 1.**
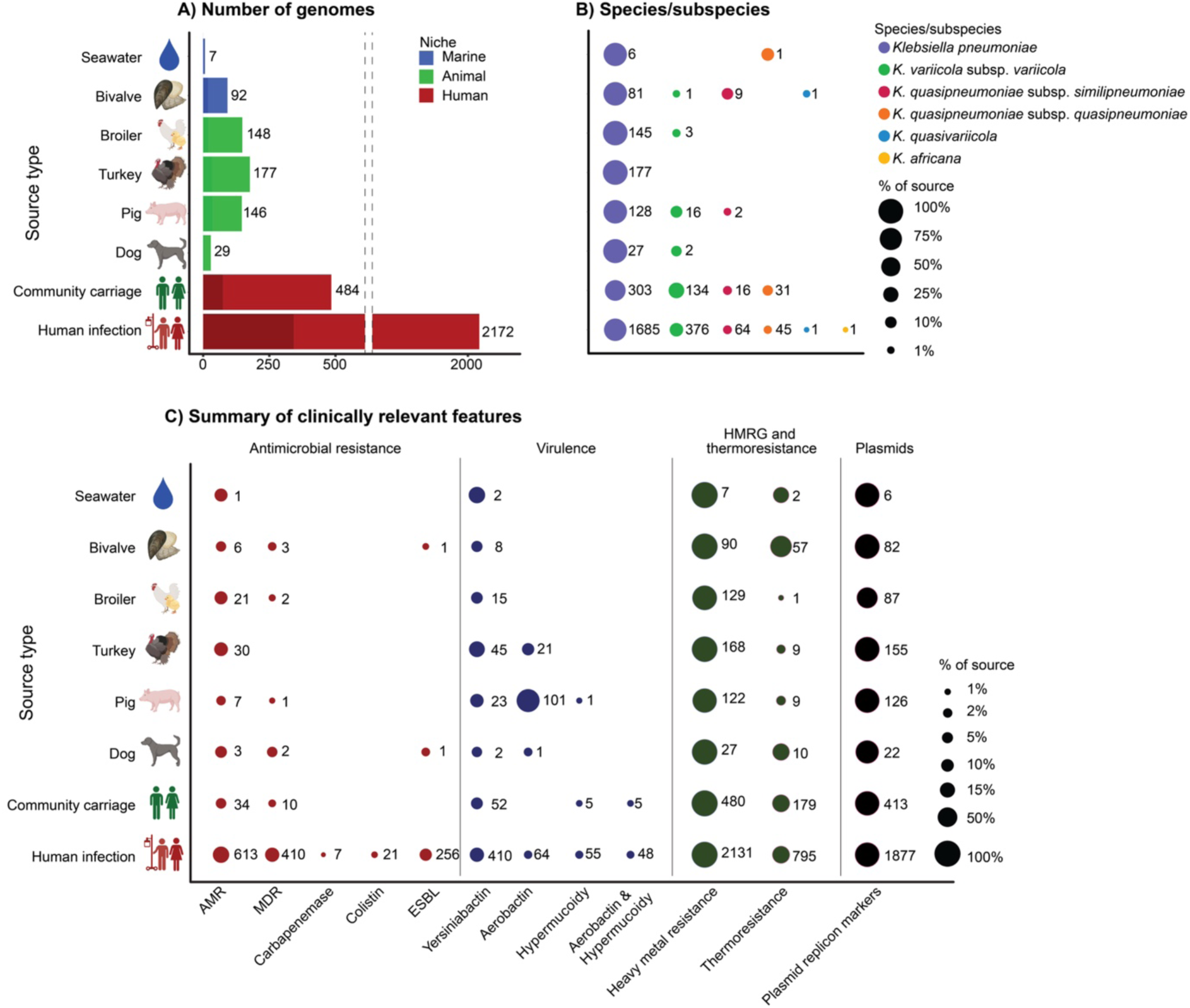
Key characteristics of the 3,255 *Klebsiella pneumoniae* species complex (KpSC) isolates. **A)** Distribution of genomes by source, with bars coloured by niche (inset legend), and shaded to indicate sequencing type (lighter for short-read only, darker for short- and long-read). Note that the x-axis is broken due to the large number of genomes from human infections. **B)** KpSC species (inset legend) distribution by source. **C)** Summary of clinically relevant features, showing the presence (bubble) of antimicrobial resistance (AMR), virulence determinants, heavy metal (HMRG)- and thermoresistance, and plasmid replicon markers across the sources. In B) and C), the bubble size corresponds to the percentage of isolates with these features, with the number of isolates shown beside each bubble.

**Table 1.** Klebsiella pneumoniae species complex carriage in Norway.

|  | Collection years | KpSC positive isolates (%) / number of samples * |
| --- | --- | --- |
| **Human niche** |  |  |
| Community carriage <sup>20</sup> | 2015, 2016 | 484 (16.3%) / 2975 |
| **Animal niche** |  |  |
| Turkey flocks <sup>21,59</sup> | 2018, 2020 | 191 (69.2%) / 276 |
| Broiler flocks <sup>21,59</sup> | 2018, 2020 | 158 (25.2%) / 627 |
| Pigs <sup>15</sup> | 2019 | 146 (49.0%) / 298 |
| Dogs | 2019 | 16 (6.5%) / 246 |
| Cattle | 2019 | 12 (4.0%) / 299 |
| Wild boars | 2020 | 27 (21.4%) / 126 |
| **Marine niche** |  |  |
| Bivalves <sup>22</sup> | 2016, 2019, 2020 | 92 (15.9%) / 578 |
| Surface seawater <sup>22</sup> | 2019 | 7 (41.2%) / 17 |
| Sea urchin | 2019 | 0 (0%) / 7 |
| Fish | 2019, 2020 | 0 (0%) / 53 |
| Sediment | 2019 | 0 (0%) / 24 |
| **Total** |  | 1134 (20.5%) / 5527 |

\* Some of the animal samples were pooled before assessing KpSC presence; see the Supplementary methods for sampling details. References are provided in the table for the sources whose carriage rates have previously been reported.

*K. pneumoniae* is the major species within the KpSC in clinical settings ^1^. In our collection, it was the most common species within all sources ranging from 62.6 to 100% (Fig. 1B). The 3,255 genomes belonged to 857 SLs (Table S1), the majority of which were represented by only a few genomes each: 87.8% (752/857) by ≤5 genomes; 59.0% (506/857) were singletons. Simpson diversity indices of SLs were 0.99 for the human, 0.98 for animal and 0.96 for marine niche. Despite high diversity, there was considerable overlap between the niches: 12.5% (107/857) were found in at least two niches, and accounted for nearly half (48.5%, 1,578/3,255) the genomes. These SLs belonged to *K. pneumoniae* (n=91), *K. variicola* subsp. *variicola* (n=12) and *K. quasipneumoniae* subsp. *similipneumoniae* (n=4).

### AMR was concentrated in human KpSC isolates

We observed low levels of acquired AMR, reflecting the low use of antibiotics in Norway in humans and animals ^24^. Still, AMR determinants were much more common in humans (23.6% with acquired AMR, 15.8% MDR, 9.6% extended-spectrum β-lactamase [ESBL]) than animals (12.2% AMR, 1.0% MDR, 0.2% ESBL) and marine sources (7.1% AMR, 3.0% MDR, 1.0% ESBL) (P <0.001, see Table S2, Figs. 1C, S1 and S2). Only two isolates from non-human sources encoded ESBLs: ST1035 with *bla*_CTX-M-3_ (from a bivalve), and ST1583 with *bla*_CTX-M-15_ (from a canine infection). Neither the strains nor plasmids from these isolates were present elsewhere in our collection.

AMR determinants were distributed across SLs, found among 24.4% (189/776) of SLs in isolates from humans, 16.2% (24/148) in animals and 13.5% (7/52) in the marine niche. Of eight recognised MDR-associated clones (SL101, SL147, SL15, SL258, SL29, SL307, SL37 and SL17)^1^ all were found among the human infection isolates. SL17, SL37 and SL29 were also seen in community carriage, animal and marine isolates, and one SL258 isolate in community carriage. Of 488 genomes with these eight SLs, only 34.8% (170) were MDR and 25.8% (126) encoded ESBLs or carbapenemases. The isolates in these SLs in the community carriage, animal and marine isolates were not MDR (except one marine SL37) and did not encode any ESBLs or carbapenemases (Fig. S3A).

### Animals as reservoirs for virulence factors

Known acquired virulence factors were not as concentrated in the human niche as AMR determinants (Figs. 1C, S2, and S4). There were no significant differences in yersiniabactin prevalence across the niches, ranging from 6.9 to 28.6% per source. Aerobactin-encoding loci were present in 6.0% (194/3,255) of genomes, the majority in animals: 10.8% (21/194) belonged to a clonal expansion of an SL290 strain that carried an IncFIB/IncFII plasmid encoding *iuc*5 and *iro*5 in turkey isolates, described previously ^21^; neither the chromosome nor plasmid were seen elsewhere in our dataset. Accounting for 52.1% of the aerobactin-encoding genomes, IncFIB(K)/IncFII *iuc*3-encoding plasmids (and one chromosome) were seen in 101 pig isolates, in 50 SLs, as previously reported ^15^*. Iuc*3 was rare in human isolates (8 infections, 2 carriage). A comparison with publicly available *iuc*3-encoding genomes^16^ showed that two of these had close relatives with the Norwegian pig isolates: one carriage-pig pair was clonally related (SL10332, 0 *iuc*3 plasmid single nucleotide polymorphisms (SNPs), 100% replicon sequence coverage), the other pair belonged to SL35 (human blood) and SL10334 (pig) and shared 4 *iuc*3 plasmid SNPs and 99.9% *iuc*3 plasmid coverage, consistent with local acquisition from pigs (e.g. via food or direct contact with pigs). The remaining *iuc*3-encoding plasmids in humans had no local relatives amongst human or pig isolates in our collection (>20 SNPs), but were related to human clinical *iuc*3 isolates from Asia or Europe (closest pairwise relatives shared 1-46 SNPs, median 12.5), as previously noted ^16^, which may reflect imported strains (e.g. via imported food or human travel).

Hypervirulence-associated clones^1^ were detected only in the human niche (SL23, SL25, SL380, SL66 and SL86), except for one SL25 genome (*ybt*6; ICE*Kp5*, K locus KL2 and MDR) in the marine niche which was closely related to an isolate from the human niche (see Fig. S3B and below). The genomes belonging to these clones had capsule locus KL1 or KL2, and carried aerobactin loci *iuc1* or *iuc*2, salmochelin *iro*1 or *iro*2 and the hypermucoidy loci *rmp*1 or *rmp*2 (except 44/45 SL25 genomes that encoded no *iuc*, *iro* or *rmp* loci).

### Heavy metal and thermoresistance associated with human and marine sources

Heavy metal and thermoresistance loci, which may facilitate survival and/or maintenance of antibiotic resistance mechanisms, were present in all niches, but the frequencies of different operons varied significantly between niches (Fig. 2). The most common heavy metal resistances for all niches were to chromium and cobalt/nickel (typically chromosomally encoded; Fig. S5 and S6). However, these showed distinct niche distributions, with *chrA/B1* (chromium resistance) significantly less prevalent in animal isolates (45.4% vs 68.5% in humans, P<0.001) and *rcnAR* (cobalt/nickel resistance) less prevalent in human isolates (65.1% vs 77% in animals, P<0.001). Silver (*sil*), copper (*pco*), arsenic (*ars*) and thermoresistance (*clpK*, *hsp20*) operons were also more prevalent in human and marine isolates (>25%) compared to animals (≤15%; P<0.001, see Fig. 2), and were typically plasmid-encoded (Fig. S5). We note a correlation between the two most common plasmid replicon markers (IncFIB(K) and IncFII(pKP91) and genes encoding resistance to several antibiotic classes (aminoglycosides, sulfonamides, trimethoprim, ESBLs) as well as heavy metals (*sil*, *pco*) and thermoresistance (Fig. S7). In total, 93 genomes from the human niche carried all of these determinants (Fig. S8), together with IncFIB and IncFII plasmid markers. They belonged to 28 SLs, the most common were SL307 (n=36), SL15 (n=7), SL45 (n=7), SL17 (n=5), SL258 (n=5) and SL35 (n=5). Additionally, several genomes in all three niches carried these plasmid replicon markers, heavy metal and thermoresistance genes without the AMR determinants.

**Fig. 2.**
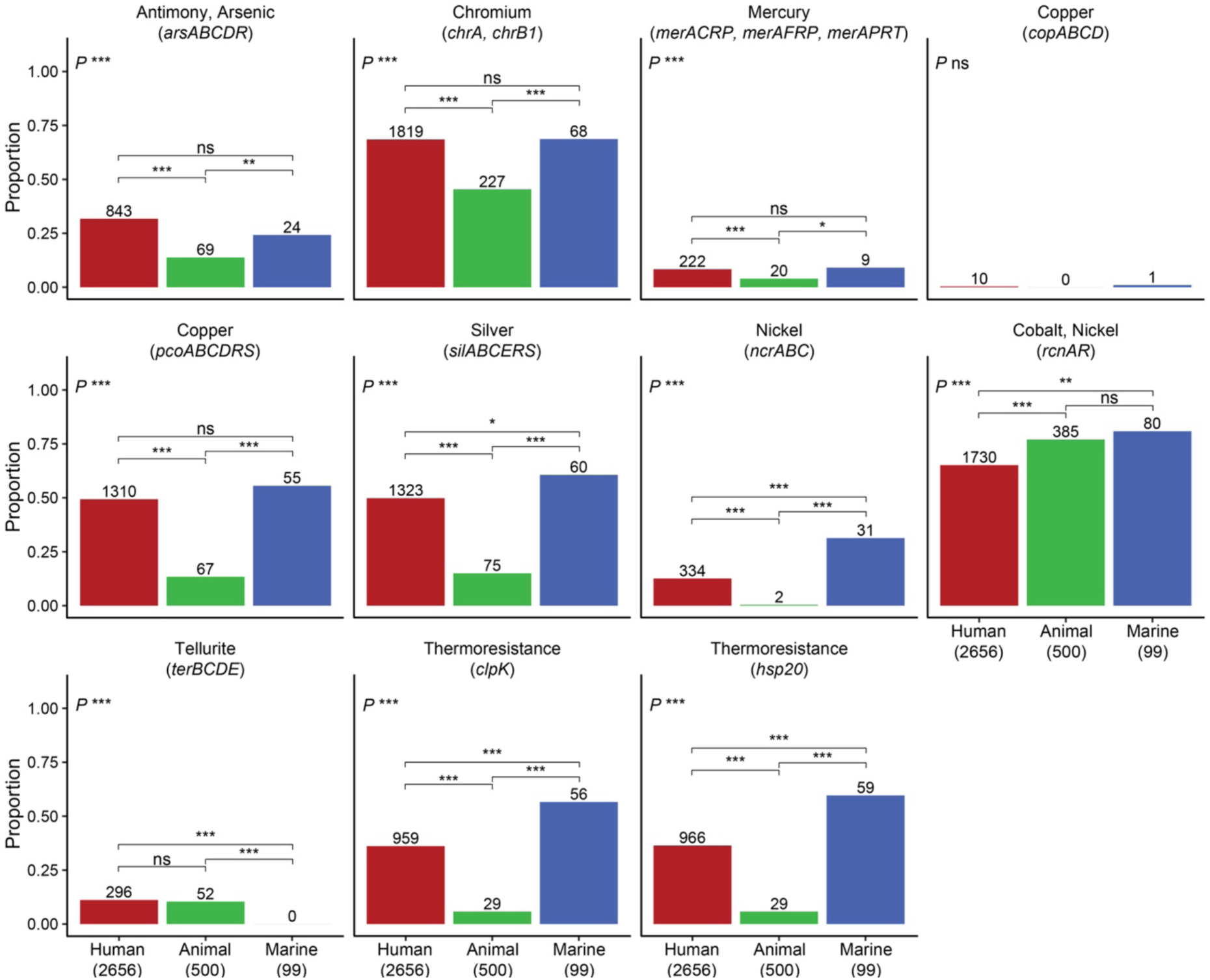
Heavy metal- and thermoresistance by niche. The proportion of genomes in the human, animal and marine niches that carried heavy metal resistance operons and thermoresistance genes. Statistical comparisons were performed with chi-squared tests. Significance levels are indicated in the plot as follows: * P<0.05, ** P<0.01, *** P<0.001, ns P≥0.05.

### Large pangenome overlap with few niche-enriched traits

We next investigated the pangenome of our genome collection and found that half the genes (53.2%; 23,748/44,614) were niche-overlapping. The human niche had a much larger accessory genome, which was expected due to the larger sample size and collection time. However, the gene content diversity and gain/loss rates were similar across the niches, as estimated by comparing Jaccard distances and gene gain/loss rates (Fig. 3).

**Fig 3.**
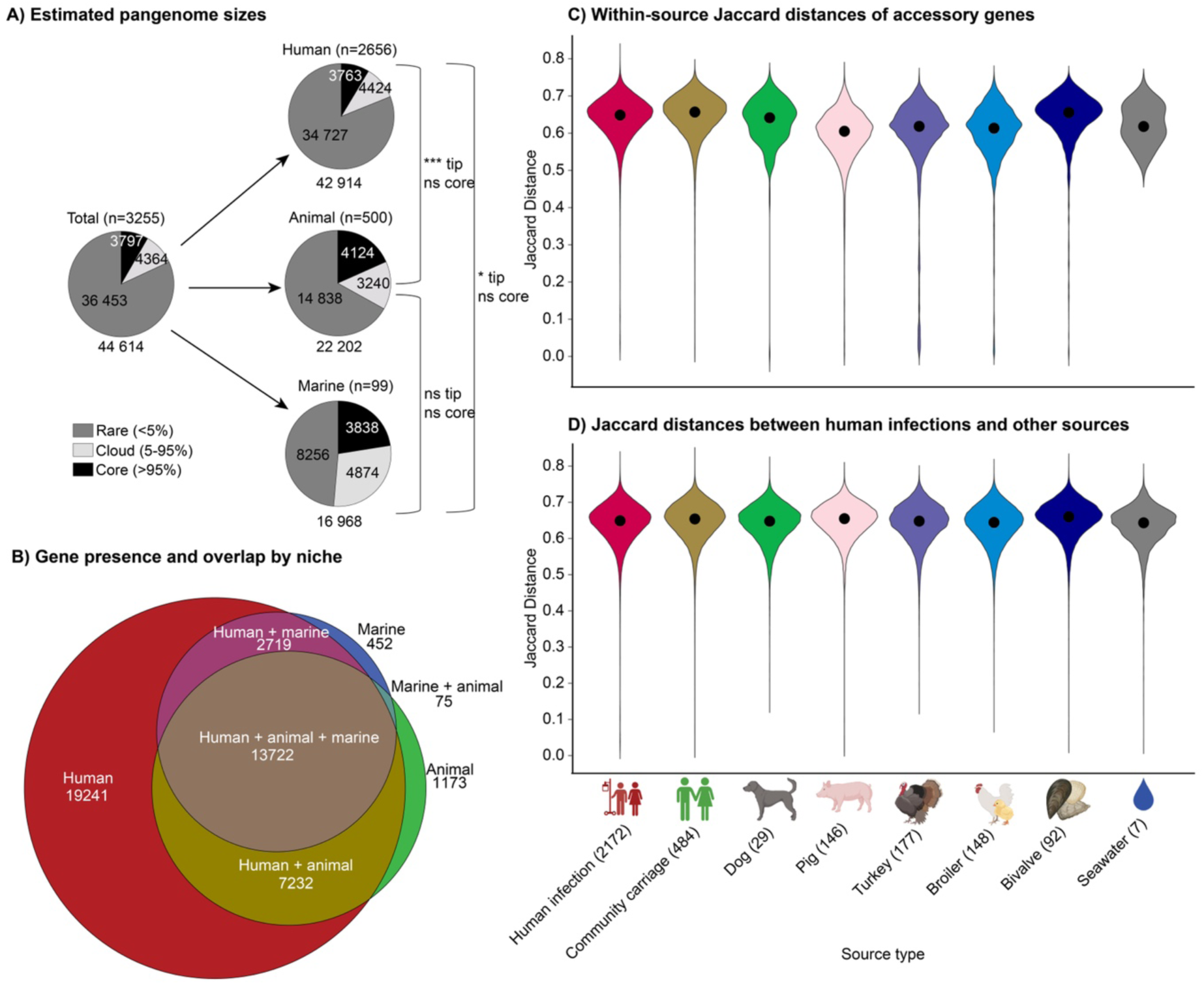
Comparison of the pangenome across niches. **A)** Overall counts of core and accessory genes, in total and by niche. A panstripe analysis indicated no significant differences in gene gain or loss rates across the niches, but the human niche had a higher rate of rare genes observed at the tips of the phylogeny (tip; P<0.001 when compared with animal genomes, and P<0.05 compared with marine genomes), which could be driven by a combination of an increased number of highly mobile or singleton genes present in the human niche in addition to technical variation of the annotation algorithm. **B)** Euler diagram showing presence and overlap of the 44,614 unique genes in the pangenome across the niches. **C)** Pairwise Jaccard distances of accessory genes within each source. Black points indicate median values. **D)** Between-source pairwise Jaccard distances of accessory genes. For each source, the Jaccard distance of each genome was compared with the distance in human infection genomes to compare diversity between sources.

We performed genome-wide association studies (GWAS) to look for genetic features of the 2,465 *K. pneumoniae* isolates associated with the animal (n=477) or human (n=1988) niches. The marine niche and other KpSC members could not be assessed due to low sample sizes. Overall, 43 genetic features were positively associated with the animal niche and 39/43 (90.7%) were also negatively associated with the human niche: 22 genes (of 36,771 unique among the *K. pneumoniae* genomes), 5 structural genes (of 54,261; i.e., groups of three consecutive genes), 16 unitigs (of 5,688,601), and no SNPs (of 959,726) (Table S3). These genetic features were enriched in animals, but were also present at low frequencies in some isolates from humans (Fig. 4), indicating that strains, MGEs or plasmids with these features in the human niche may have originated from the animal niche or have become associated with the animal niche once introduced there (we could not infer directionality).

**Fig. 4.**
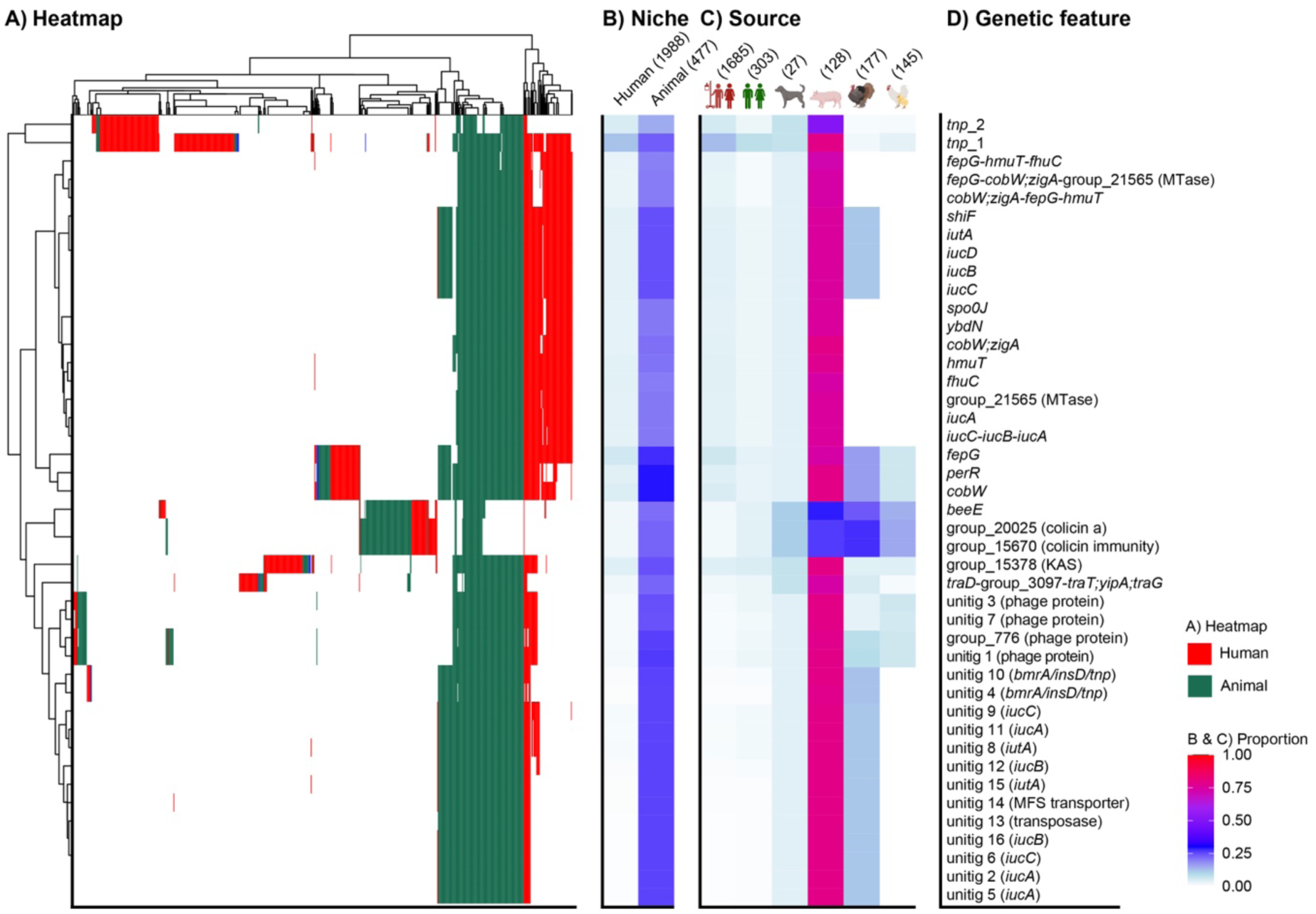
Niche-associated genetic features identified through genome-wide association studies (GWAS). **A)** A heatmap showing the presence of significantly associated genetic features (rows), among the 2,456 *Klebsiella pneumoniae* genomes from humans and animals (columns), coloured by niche (inset legend). All features were positively associated with the animal niche. **B)** Proportions of genomes with genetic features by niche and **C)** source. The total number of genomes per niche and source are indicated on top of the columns. **D)** Annotation of genetic features. See Table S3 and Fig. S9 for more detail. Genes annotated with more than one name are separated by a semicolon, unitigs found in multiple genes are separated by a slash.

Many of the features were linked to the aerobactin *iuc* locus, which has previously been identified as pig-associated (see above) ^15,16^. Most of the genes and unitigs were co-distributed with *iuc* genes and similarly associated with animals in the GWAS, presumably due to physical linkage on the *iuc*-encoding plasmids (Fig. S9A), although they were not all exclusively found in *iuc*-encoding genomes (see Fig. S9B). These features were likely enriched in the animal niche by the increased prevalence of the pig-associated *iuc*3-encoding plasmids. The genes enriched in the animal niche not associated with *iuc* plasmids were two encoding the bacteriocin colicin a and colicin immunity protein (group_20025 and group_15670; see Table S3), and a phage portal protein (*beeE*), which were frequently found together on plasmids, although *beeE* was also found separately on several chromosomes (Fig. S9B). The colicin genes were found in 27.7% (49/177) of *K. pneumoniae* isolates from turkeys, 26.6% (34/128) pigs, 15.2% (22/145) broilers, and 11.1% (3/27) dogs, compared to 1.6% (27/1,685) human infection and 3.6% (11/303) community carriage. There was no clear association with particular clones, except for 15 genomes belonging to SL35 with KL22. Inspection of 14 closed genomes with these features revealed that colicin a, colicin immunity protein and the phage protein were located on highly similar plasmids (Fig. S10).

### Lack of niche restriction of KpSC sublineages

Some SLs were broadly distributed across sources: SL17 (n=177), SL37 (n=144), SL45 (n=81), SL200 (n=34), SL29 (n=32), SL34 (n=30) and SL111 (n=21) were identified in all niches (Table S1). SL107 (n=103), SL35 (n=78), SL641 (n=36) and SL290 (n=35) were found in humans and animals but not marine samples, and SL3010 (n=123), SL10 (n=48), SL25 (n=45) and SL461 (n=33) in humans and marine samples but not animals. The detection of these SLs across niches cannot be interpreted as direct transmission between niches, however it argues against niche restriction of SLs. Dated phylogenies showed that these clones have existed for a long time: estimated dates for the most recent common ancestors (MRCAs) of genomes of each SL from Norway ranged from 1313 to 1972, with variable uncertainty (Fig. S11 and Table S4). Further, a search of publicly available genomes at Pathogenwatch ^25^ showed that these SLs are widely distributed over time, geography and sources: 4-58 countries, 10-23 years, and at least one genome per SL was found in animal, food, environmental or wastewater samples (Table S5).

The most prevalent SLs identified in a single niche were found in humans only: SL307 (n=62), SL14 (n=46), SL1562 (n=28), SL15 (n=26), SL268 (n=25), SL2004 (n=25), SL359 (n=23), SL258 (n=22), SL70 (n=21) and SL23 (n=20). This was not surprising as the sample size for humans was much larger than for other niches. However, there was evidence to support that the prevalence of these SLs in non-human samples is actually lower: if the SLs with >23 human isolates were present in the other niches at the same prevalence as in humans, we would expect to observe them at least once amongst the animal and marine samples (assessed using a binomial test with the alternative hypothesis that they are less prevalent, see Table S6A). Therefore, these SLs may be present in the animal and marine sources, but at significantly lower frequencies than amongst human isolates.

Most capsule biosynthesis (K) (79/131, 60.3%) and lipopolysaccharide (O) loci (10/12, 83.3%) were observed across niches. However, some loci were enriched. O1/O2v2, O3/O3a, O4 and OL103 were enriched in the human niche, whilst O3b and OL13 were enriched in the animal niche (see Table S6B). Of K loci, KL102, KL103, KL2, KL10 and KL25 were enriched in the human niche; KL21, KL14, KL22, KL30, KL31, KL55 and KL57 in the non-human niche. Several of these K and O loci were associated with particular strains, e.g. SL107, KL103 and O1/O2v1 were found in 86 genomes that were part of a clonal expansion (see below and Table S6C). K and O loci were relatively stable within SLs, and most SLs were associated with a single K/O combination (82.6%; 708/857), and SLs from different sources shared the same K/O loci (Fig. S12).

### Cross-niche strain-sharing rare, but of potential impact

To assess the relatedness of strains within and between niches, we enumerated pairwise genome-wide SNPs within the 107 niche-overlapping SLs (Fig. 5A). Isolates from the same niche were more closely related (*P*<0.001, see Fig. 5B), consistent with within-niche transmission. However, we also observed closely related pairs of isolates from different niches. The SNP distances between isolates from human infections and those from other sources showed that human infection isolates shared more recent ancestry with isolates from human community carriage, followed by pigs and bivalves (Fig. 5C). At ≤22 SNPs (selected using cutpointR as the optimal threshold to differentiate within- vs. between-source in this study [see Methods], and in line with thresholds identified in hospital-based studies to identify transmission clusters ^4,26,27^), 1.9% (41/2,172) of human infection isolates were linked to community carriage isolates in our sample (which were restricted to individuals living in the Tromsø municipality, isolated 2015-2016, compared with clinical isolates which were sampled across all Norwegian hospitals during 2001-2018). At the same genetic distance (≤22 SNPs), 3.9% (86/2,172) of human infection isolates were linked to pig isolates (n=5; collected in 2019 from 5 farms across the country) and 0.8% (18/2,172) to bivalves (n=3; collected in 2016 and 2019 from different locations, two from food production sites).

**Fig. 5.**
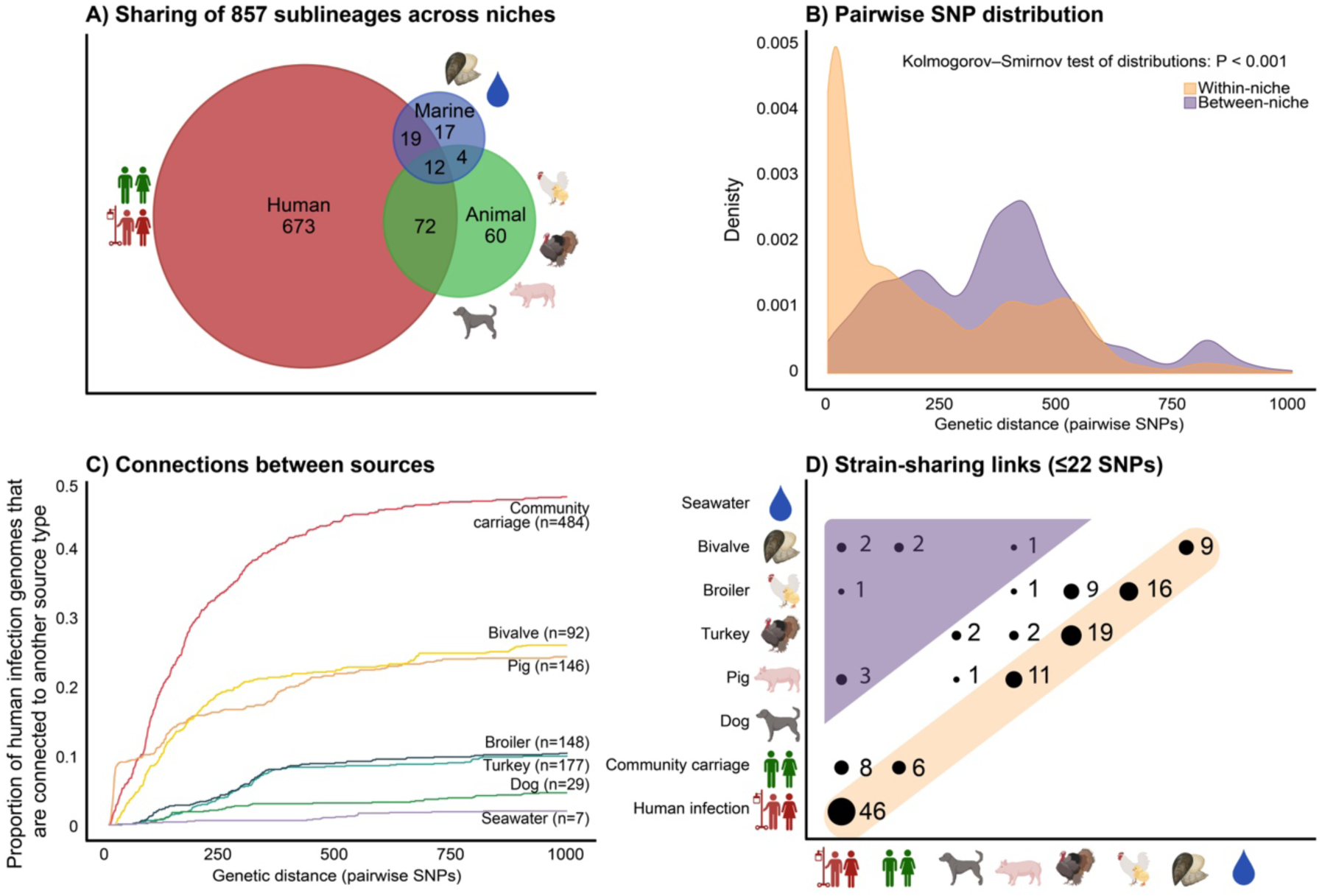
Strain-sharing within and between ecological niches and sources. **A)** Venn-diagram showing the distribution of 857 sublineages (SLs) across human, animal and marine niches. In total, 107 SLs were present in ≥2 niches. **B)** Distribution of pairwise single nucleotide polymorphism (SNP) distances, grouped by genome-pairs within the same niche (orange) and between niches (purple), showing distances up to 1000 SNPs. **C)** Connections between sources. The proportion of human infection genomes (y-axis) that were connected to one of the other sources, by the number of pairwise SNPs (x-axis). **D)** Strain-sharing links (clusters of genomes sharing ≤22 SNPs) between source pairs, as indicated on the x- and y-axes. The yellow shading highlights within-source clusters, and the purple highlights the cross-niche clusters.

Using the ≤22 SNP threshold, there were 130 within-niche sharing events (60 within the human niche, 61 animal and 9 marine), compared to only nine between the niches (Fig. 5D). Isolate pairs with ≤22 SNPs were collected within short time periods (mean 0.68 years, range 0-11). The nine cases of cross-niche strain-sharing were between humans and pigs (1 SL37, 2 SL107), humans and broilers (1 SL200), humans and bivalves (2 SL25, 1 SL461, 1 SL3010), and pigs and bivalves (1 SL3676) (Fig. S13). Dated phylogenies of SL25, SL107 and SL3010 showed that the cross-niche pairs had MRCAs <3.8, <5.46 and <14.5 years ago, respectively (Fig. S11).

## Discussion

KpSC infections in humans can be caused by strains circulating within the human population or introduced from animals or the environment as a result of contaminated food or direct contact. Using population genomics in a One Health context, we have shown that the KpSC populations isolated from human infection and community carriage, animal and marine sources from Norway are distinct but overlapping. In particular, pigs and marine bivalves appear as potential reservoirs for strains and plasmids that contribute to colonisation and infection in humans. Our study was conducted in a low AMR prevalence context. In the Norwegian surveillance programme for AMR in 2023, only 6.6% of human clinical KpSC isolates were resistant to third-generation cephalosporins and 0.4% to carbapenems ^24,28^. This provided an opportunity to investigate the interactions of KpSC populations without interference of strong selective pressures from antibiotics.

In total, we identified nine cases of recent strain-sharing between the niches. This national collection was sampled during a 20-year period, but the samples in the three niches were not matched by year or location. We therefore did not expect to capture direct transmission links. However, the detection of genetically related isolates implies shared ancestry, and thus movement of bacterial strains between niches, although we could not ascertain directionality nor confirm whether movement occurred directly between the sampled niches, via intermediate niches or from a common source. Contemporaneous sampling in a localised area would likely capture more sharing events, however, given the ubiquity and diversity of KpSC within sources it would be difficult to capture evidence of transfer without sampling directly linked sources (e.g. humans and the specific food sources they were recently exposed to).

We found that spillover of strains and clinically relevant genes was more common between sources within the same niche. This was not unexpected for the human niche, where it has been shown that as much as half of hospital-acquired KpSC infections are caused by a patient’s own colonising strain on admission ^4^. There were several strain-sharing events within and between different food-producing animals (pigs, turkeys and broilers). Strain-sharing within the same host may be due to vertical dissemination in animal production pyramids, and sharing both within and between sources may be explained by unsampled transmission from other animals or people, or from e.g. tools, feed or fertiliser ^29^. Few events were observed in dogs (only with pigs and turkeys), likely due to the small sample size and unmatched dataset. The marine niche was the most underrepresented in our collection. Yet, five of the nine cross-niche events were linked to marine bivalves, and the marine isolates were more similar to the human isolates in terms of heavy metal and thermoresistance genes compared to the animal isolates. Bivalves are filter feeders that can accumulate pathogens from their surrounding environment. Agricultural runoff or sources of human pollutants, such as wastewater effluent, are likely transmission routes from humans to bivalves ^30,31^, and they may also feed back to the human niche via food consumption ^32^.

Individual events of ecological spillover may be very rare, as our data and other recent studies of KpSC ^5,9,10,17^ suggest, although arguably none of these studies have sampled sufficiently from within direct-contact chains to measure the incidence of such events. Still, even rare events can have major consequences in the long term. It is clear that at least some strains of KpSC can transmit easily between humans, especially amongst vulnerable individuals in hospital settings ^23,33^. Therefore, upon a new strain entering the human niche (e.g. via food consumption), it is possible that it may spread, adapt and evolve depending on exposures, e.g. to other microbes by sharing of MGEs, or to selection dynamics including antibiotics. For example, we observed diverse SL107 isolates in pigs (consistent with persistent circulation in pigs over a decade, see Fig. S11), as well as a rapid clonal expansion of SL107 in humans (86 cases in 1.6 years, with <43 pairwise SNPs [median 13], MRCA <5 years and accounting for 4.5% of the human bloodstream infections). Whilst our data does not tell us about directionality of movement of SL107 between humans and pigs (the clonal expansion detected in humans was sister to, not emergent from within, the pig clade, see Fig. S11), it does demonstrate that the same strain can both persist in pigs and spread rapidly in humans. The SL107 strain was largely without AMR or virulence factors, but six human blood isolates had acquired either AMR determinants or virulence factors, likely reflecting shifting selective pressures in the human niche.

There was overall high genetic diversity throughout the dataset, but most prominent in the human niche, which accounted for 43.1% of the unique genes including a wider range of AMR and virulence determinants. This is likely reflective of the larger size of the human niche collection, and that people tend to travel and interact with other people, animals, and environments, thereby facilitating the spread of bacteria. In our previous study of human *K. pneumoniae* carriage, travel was associated with a higher prevalence of carriage ^20^. Still, the animal and marine KpSC populations greatly overlapped with those in the human niche. Whilst only 2.6% of the genes in the animal niche were unique to the niche, we did identify niche-associated traits that were positively associated with animals. They were largely linked with aerobactin-encoding plasmids which were frequent in animals, particularly pigs, but there were also genes linked to colicins and phages, which we hypothesise may reflect niche adaptation as a result of interactions with specific bacteria or phages unique to the animal microbiomes. Bacteriocins like colicin are peptides that can kill or inhibit the growth of competing species, offering an advantage to the colicin-producing bacteria for surviving in environments with limited nutrients or space ^34^.

In our collection of 857 SLs, 107 were present in at least two niches. SL17, SL35, SL37, SL45 and SL107 were the five most common niche-overlapping SLs. SL107 was inflated due to a clonal expansion among human infection isolates ^23^, whereas the others appeared more generalist, being common in all eight source types (except seawater for SL35). SL17 has previously been described as a generalist clone with a wide range of AMR and virulence determinants that can colonise and cause opportunistic infections in both humans and animals ^35^. Based on our data, similar conclusions may be drawn for SL35, SL37 and SL45, which were present across the niches, with varying levels of both AMR and virulence. Corroborating this, a hospital surveillance study from Australia found that SL17, SL35, SL37, as well as SL111, SL629 and SL661, are clones that frequently cause hospital-acquired infections that are not linked to nosocomial transmission, but rather highly prevalent in humans likely due to being animal-adapted strains that humans are exposed to via the food chain ^33^. These SLs have also been reported in animals and food products in other studies from Germany, Italy, Thailand, Ghana, Australia and the USA ^5,17,36–39^. SL107 may also be a generalist clone; SL107 has been observed encoding ESBL genes in animals in the Netherlands ^40^ and Germany ^36^, aerobactin (*iuc*3) in animals in Norway ^15^, Italy ^6^ and Germany ^36^, carbapenemases in clinical settings in China, and convergence of colistin resistance and aerobactin in a pig in China ^41^, whilst also causing a largely susceptible hospital outbreak in Norway ^23^. While some clones were more generalist, there were also potential specialist clones within our dataset; 673 SLs were present only in the human niche. The most abundant of these were SL14, SL15, SL23, SL70, SL147, SL258 and SL307, which have all been described previously as MDR- or hypervirulence-associated clones that disproportionately contribute to the global burden of hospital-associated disease and nosocomial outbreaks ^1^. In a recent study of bloodstream infections included in our dataset, a higher 30-day mortality rate was associated with infections caused by *K. variicola* or global MDR-associated clones (all *K. pneumoniae*) compared to other KpSC lineages ^23^. Of global MDR clones, we observed only SL17, SL29 and SL37 outside of human infection isolates (in both community carriage and non-human sources), and of *K. variicola* SLs, 5.3% (12/225) were found across niches. None of the *K. variicola* strains we observed outside of human infections were MDR (except one SL697 community carriage), and in fact most human clinical isolates of *K. variicola* were not MDR, hinting to other factors beyond AMR driving their presence amongst clinical isolates ^23^.

Another important aspect of One Health is the transfer of MGEs between strains and across niches ^42^. This will be addressed in detail elsewhere. In this study, we focused on the mobility of known clinically relevant features, including virulence and resistance to third-generation cephalosporins and carbapenems. The majority of AMR spread occurred within the human population. Genes encoding ESBLs or carbapenemases were nearly exclusively in the human niche, except for two ESBL-encoding plasmids identified in a dog and a marine bivalve. The two were not related to other strains or plasmids in our dataset, however, a KpSC strain with a *bla*_CTX-M-15_-encoding plasmid in the marine bivalve was highly similar to one recovered from a human bloodstream infection in South Korea one year later ^22^, indicating undetected transmission of some kind. We observed a correlation between the presence of AMR, heavy metal and thermoresistance genes with specific plasmid replicon markers in the human niche and found the same feature combinations without AMR present in all niches. Further data and analysis are needed to understand if these are the same or similar plasmids with/without the AMR encoding genes. A previous study hypothesised that exposure to antibiotics is the major driver behind co-occurrence of AMR and heavy metals in bacteria from humans and domestic animals ^43^. We detected a lower prevalence of MDR and heavy metal resistance determinants among animal isolates, particularly of plasmid-encoded arsenic, copper and silver-resistance genes, which may play a role in limiting innate immune responses ^44,45^. In contrast, a recent Portuguese study observed that >60% of poultry KpSC strains from several farms were MDR and resistant to copper and silver with genetic similarities to isolates from human infections ^46^, showing the potential for animals to be reservoirs for these features if they are exposed.

Known virulence-encoding genes were most frequent in the animal niche, but consisted of only two lineages: 1) a clonal expansion with an aerobactin (*iuc*5) +/- salmochelin (*iro*5)-encoding plasmid observed only in turkeys and in no other sources ^21^, and 2) the pig-associated aerobactin lineage *iuc3*, which we observed in pigs^15^, a dog, human infections and community carriage. Global epidemiological analyses of the *iuc*3 lineage previously revealed that most of the *iuc*3-encoding plasmids recovered from humans were related to plasmids originating from Asia or Europe, but two of the plasmids were likely the result of zoonotic spillover of strains between pigs and humans in Norway ^15,16^. Several reports have shown the emergence of convergent KpSC strains carrying both AMR genes and hypervirulence traits ^1,47^. We observed only six such cases, but the recent findings of plasmids harbouring both *iuc*3 and ESBLs in Thailand in both humans and animals ^16^ calls for concern given the high prevalence of *iuc*3 in our dataset.

Our overall findings are similar to KpSC One Health studies from Italy, and confirm on a much broader scale those from the UK, the USA, the Caribbean and Ghana ^5,9,10,17,18^. Human-to-human strain-sharing in clinical settings was more common than cross-niche, and resistance to third-generation cephalosporins, colistin and carbapenems were largely confined to human clinical settings. These studies suggested that infection prevention measures should be focused on clinical settings to limit the spread of MDR and to break nosocomial transmission chains. While we agree, we also emphasise that a One Health surveillance framework remains important. KpSC classically causes opportunistic nosocomial infections, however, hypervirulent and sporadic classic infections also occur in healthy individuals in the community, which cannot be prevented by hospital infection control measures ^1^. Further, it was recently shown in Norway that over half of 1,078 KpSC bacteraemia cases were community acquired as opposed to hospital or healthcare-associated ^23^. Infection is often preceded by colonisation as a result of human-to-human transmission, or direct contact with animals or contaminated food. Therefore, other interventions that prevent or limit transmission and subsequent colonisation from the food chain are also important. Though rare, the observation of cross-niche transmission despite Norway’s rigorous livestock management and biosecurity measures highlights the continued relevance of One Health approaches also in high-income countries ^29^.

Ecological barriers such as geographical distance or physical barriers are likely limiting exposure (opportunity for frequent strain transmission) between niches more than the genetic make-up of the KpSC themselves. We have here shown that over half the genes in the pangenome overlap the three ecological niches, and a recent study also found considerable overlap of accessory genes between the community carriage isolates in Norway and isolates from Ghana ^17^. One Health studies of KpSC to date have found little overlap of AMR between humans and animals ^5,9,10,17,18^, however, whilst the prevalence may be low, MDR and/or ESBL KpSC strains have been found in animals, vegetables and animal food products from several geographical regions, with similarities to clinical strains, including global MDR-associated clones ^18,46,48–54^. A study performed in six countries in Europe found little overlap of AMR between food items and human-polluted environments ^55^. In contrast, a systematic review of 38 studies from low- and middle-income countries showed that food-producing animals are important reservoirs for clinically relevant resistance determinants and urged that increased surveillance is necessary for aiding prevention in a similar manner to that developed by the USA and countries in the European Union ^56^. Another study found that AMR-encoding bacteria were distributed among humans, animals and the environment in Tanzania ^57^, noting that the amount of direct or indirect contact between food-animal reservoirs and people affect the amount of transmission, and that this varies based on socio-economic, cultural and ecological contexts. Thus, the low rates of AMR, virulence and strain transmission across niches in our study and in other high-income countries likely reflect the efficiency of existing preventative measures, highlighting the success of One Health initiatives rather than diminishing their importance.

To conclude, our findings together with One Health studies in other settings indicate that human-to-human transmission of bacterial strains and AMR is more frequent than between ecological niches, and even more so within clinical settings. However, our study also identified the animal niche as a reservoir for virulent strains that may spillover to humans and cause highly pathogenic infections. Taken together with growing evidence of convergent plasmids also in non-human niches, these observations reinforce the notion that the potential for public health implications from even rare zoonotic transmission must not be underestimated. Our findings support the case for surveillance of bacteria in a One Health perspective even in a low AMR setting like Norway, as it enables detection of emergence and spread of resistant or pathogenic strains that could have significant public health impacts. Increased population densities and environmental changes will likely lead to more frequent interactions between human, animal and environmental niches, which will facilitate zoonotic transmission and the formation of new reservoirs, underscoring the need for One Health surveillance approaches globally.

## Materials and methods

### Sample selection and whole-genome sequencing

We included 3,255 KpSC isolates in this study. They were collected between 2001 and 2020 from human infections (n=1,920 blood, 252 urine) ^19,23^, human faecal carriage (n=484) ^20^, from the marine environment (n=92 bivalve molluscs, 7 surface seawater) ^22,58^ and from animal caecal or faecal carriage or infections (n=177 turkey and 148 broiler flocks, 146 pigs, 29 dogs) ^15,59^ (details in Supplementary Methods). The isolates were Illumina short-read sequenced using the Illumina MiSeq (n=3,218) or HiSeq 2500 (n=37) platforms as described previously ^19–21,58^, adapter- and quality filtered with TrimGalore v0.6.7 (https://github.com/FelixKrueger/TrimGalore) and assembled with SPAdes v3.15.4 ^60^. We additionally performed Oxford Nanopore Technologies long-read sequencing of 16.9% (n=550/3,255) of the collection, to produce closed hybrid genomes, as described in ^61^.

### Genotyping and annotation

Kleborate v2.4.0 ^62^ was used to identify species, STs, virulence and AMR genes. K and O loci were identified with Kaptive v2.0.8 ^63^. The presence of the *rmpADC* locus (with/without truncated *rmpA*) was used to define a likely hypermucoid phenotype ^64^. Plasmid replicon markers were identified with plasmidfinder-db v2023-01-18 (https://bitbucket.org/genomicepidemiology/plasmidfinder_db.git) using abricate v1.0.1 (https://github.com/tseemann/abricate). Life identification number (LIN) codes based on cgMLST profiles were assigned to define and name KpSC SLs according to the 3-level LIN code prefixes ^65^, using the *Klebsiella* BIGSdb-Pasteur web tool (https://bigsdb.pasteur.fr/klebsiella/). SLs were compared to globally prevalent clonal groups (CGs) ^1^ with the same number, except for SL17 which was considered equivalent to CG20. Bakta v1.8.1 ^66^ with database v5.0 was used to annotate all assembled genomes, using the complete flag for circularised hybrid-assembled genomes and otherwise default settings. Bakta was run with NCBI’s AMRFinder database v2023-04-17.1, from which heavy metal and thermoresistance genes were identified (see Supplementary Methods). The genotyping results are available in Table S7.

### Assessing niche enrichment

To assess differences in genetic diversity between the niches, and enrichment of genetic features in the three ecological niches, we first estimated the pangenome with panaroo v1.3.3 using the strict mode ^67^, using the annotations from Bakta as input. Panstripe v0.1.0 ^68^ was then used to compare gene gain and loss rates between the niches. To assess niche-enrichment of genes, unitigs and SNPs, we performed GWAS with pyseer v1.3.11 ^69^ as described in the Supplementary Methods.

### Inferring cross-talk between niches

There were 107 defined SLs represented by ≥1 genome in ≥2 ecological niches. To assess the relatedness of strains within these SLs across niches, we created reference genomes for each and ran whole-genome SNP alignments using RedDog v1beta.11 (https://github.com/katholt/RedDog), as described previously ^35^, aligning all short-read genomes belonging to that SL against the reference genome (overall mean chromosomal coverage was 96.9% [range 88.6-100%], Table S1). Pairwise SNP distances were identified from the alignments with snp-dists v0.8.2 (https://github.com/tseemann/snp-dists). We identified an optimal cutoff-point of ≤22 SNPs using cutpointR v1.1.2 (https://github.com/Thie1e/cutpointr, see Fig. S14). To record strain-sharing events, networks were created using a similar method as in ^5^: using a SNP threshold of ≤22 SNPs, and counting each source-pair (including same-source pairs) only once per SNP-linked cluster. We also inferred dated phylogenies of SLs that had ≥20 genomes and were sampled over at least a 5 year-period, to ensure sufficient temporal signal (Table S4 and S5), in total 15 SLs, of which seven converged and had significant clock signal. Verticall distance v0.4.2 (https://github.com/rrwick/Verticall) was used to filter recombinations and BactDate v1.1.1 ^70^ to infer the phylogenies (see Supplementary Methods).

### Definitions

Multidrug resistance (MDR) was defined based on presence of AMR genes or mutations identified by Kleborate: Kleborate num_resistance_classes count ≥3.

### Statistical analyses

Statistical analyses were performed with R version 4.4.0 (2024-04-24). Comparisons of proportions were analysed with Chi-squared tests, ranges with Kruskal-Wallis test (for overall groups) and Mann-Whitney U test (for each pair of groups), and distributions with Kolmogorov–Smirnov test. Binomial tests with Bonferroni correction were used to test whether features found in humans or non-human sources were overrepresented in the other. All tests were two-tailed. P-values <0.05 from these tests were considered statistically significant, and significance was reported as follows: * P<0.05, ** P<0.01, *** P<0.001, ns P≥0.05. Significance thresholds for the GWAS results were decided by inspecting QQ-plots (Fig. S15).

## Data availability

All Illumina and Oxford Nanopore Technologies reads are available under the umbrella project PRJEB74192 on the European Nucleotide Archive. The hybrid-assembled genomes have been deposited in GenBank. See Table S8 for read and biosample accessions and metadata. The code used for analyses and to create figures is available in: https://github.com/marithetland/KGWP1_crossniche.

## Acknowledgements

We wish to thank the Norwegian Study Group on *Klebsiella pneumoniae*, the NOR-KLEB-NET and KLEB-GAP partners, the Institute of Marine Research in Norway, the Norwegian Veterinary Institute, the Norwegian National Advisory Unit on Detection of Antimicrobial Resistance, the Tromsø7 population survey, and the NORM/NORM-VET programs for their efforts in collecting and/or providing samples of KpSC, that were whole-genome sequenced at the Department of Medical Microbiology, Stavanger University Hospital, Norway. We thank the Institut Pasteur teams for the curation and maintenance of BIGSdb-Pasteur databases at http://bigsdb.pasteur.fr/. We also thank Ed Feil, Erkison Odih, Zoe Dyson, Hassan Al-Mana, Ingvild Dalen, Anna Bjørheim and Benjamin Silvester for valuable discussions. The source icons in Figs. 1, 3-5, S1, S3, S4, S6, S10 and S12 were created with BioRender.com (Created in BioRender. Hetland, M. [2024] BioRender.com/x91j569).

## Author contributions

Conceptualization: MAKH, SB, BTL, ØS, MS, AS, AMU, IHL, KEH. Methodology: MAKH, KEH. Software: MAKH, MAW, MMCL, RRW, KEH. Formal analysis: MAKH, MAW, HK, AC, RRW, KEH. Investigation: RJB, EB. Resources: HK, FH, AF, BTL, NPM, NR, ØS, MS, AS, AMU, IHL. Data Curation: MAKH, MAW, HK, FH, MMCL, JFD, SB. Writing - Original Draft: MAKH, MAW. Visualisation: MAKH, MAW. Supervision: ØS, IHL, KEH. Project administration: MAKH, IHL. Funding acquisition: MAKH, SB, BTL, ØS, MS, AS, AMU, IHL, KEH. All authors contributed to data interpretation, reviewed and edited the manuscript, and all authors have read and agreed to the published version of the manuscript.

## Ethics declarations

No ethical approval was needed for this study.

## Abbreviations

AMR: antimicrobial resistance
CG: clonal group
ESBL: extended-spectrum β-lactamase
GWAS: genome-wide association studies
KpSC: *Klebsiella pneumoniae* species complex
MDR: multidrug resistant
MGE: mobile genetic element
MRCA: most recent common ancestor
SL: sublineage
SNP: single nucleotide polymorphism
ST: sequence type

## Supplementary methods

Please see separate PDF for the Supplementary Methods.

Please see separate Excel file for the Supplementary Tables:

## Supplementary tables

**Table S1**. Source-overlap by sublineage

**Table S2.** Chi-squared tests of AMR presence by niche

**Table S3.** GWAS results

**Table S4.** BactDating results

**Table S5.** Genomes included in BactDating analyses

**Table S6.** Binomial tests of SLs, KLs and OLs by human vs non-human sources

**Table S7.** Genotyping results

**Table S8.** Genome accessions and metadata

---

**Fig. S1.**
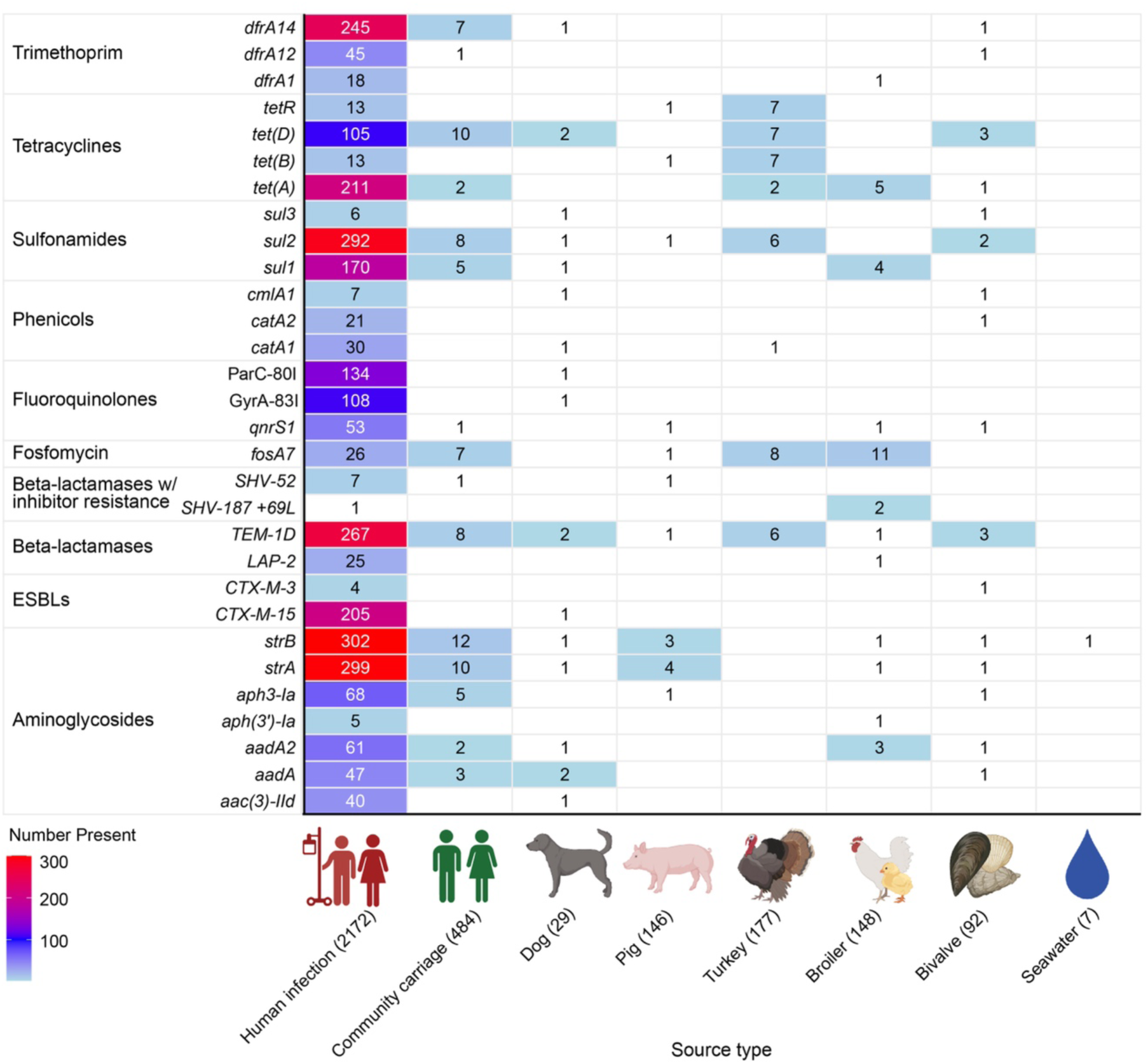
Antimicrobial (AMR) genes and mutations by source. The number of genomes per source (columns) that have the AMR gene or mutation (determinant) specified in the rows. The AMR determinants are ordered by the AMR class they belong to. In total, 130 AMR determinants spanning 14 AMR classes were observed, of which 127 were found among human infections. Here, only AMR determinants that were also found in ≥1 non-human isolate are shown (n=42). The determinants were identified with Kleborate, and includes all complete hits and those with mismatching nucleotides or incomplete (>90%) coverage.

**Fig. S2.**
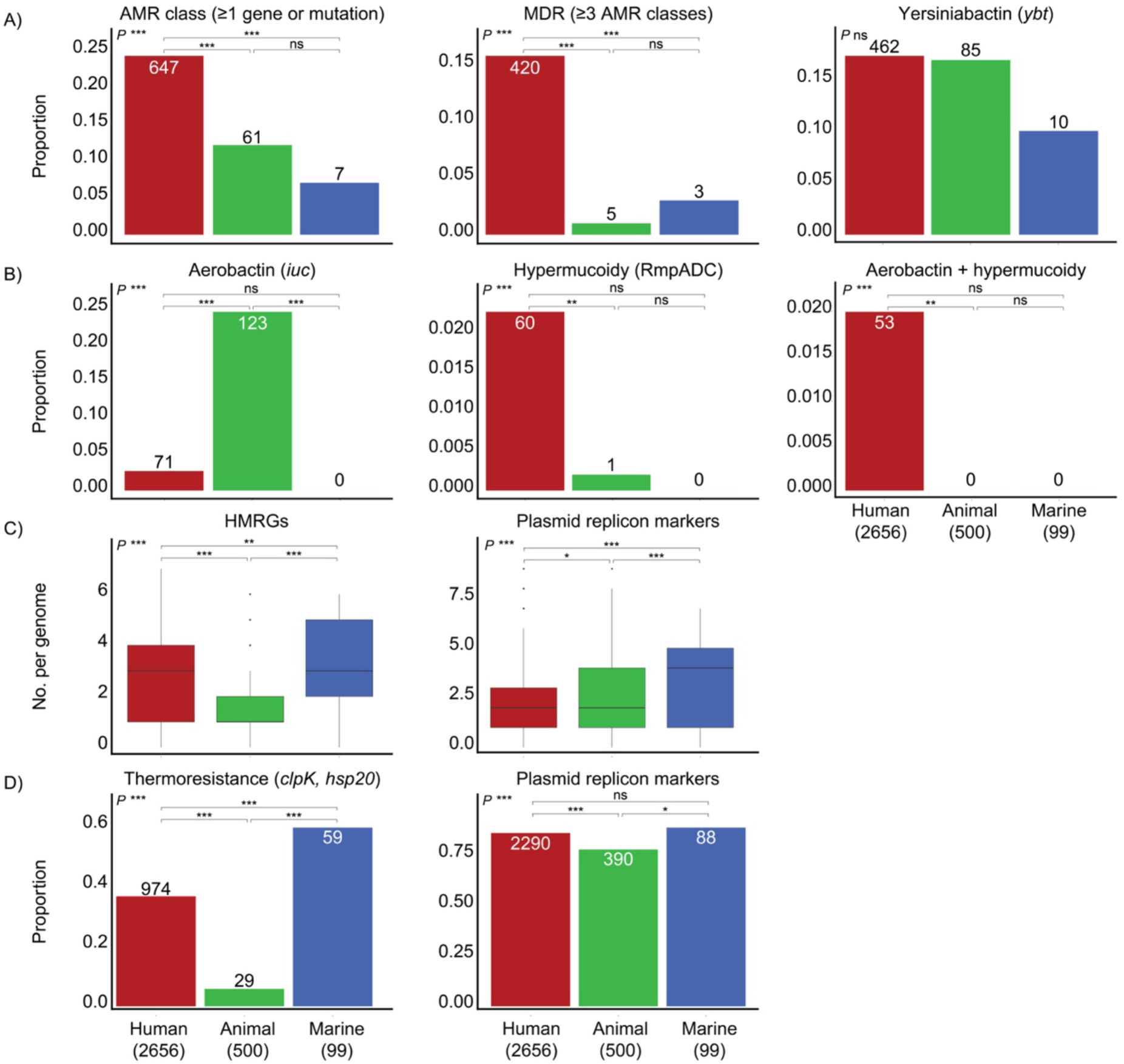
Comparison of clinically relevant features across niches. **A and B)** Proportion of antimicrobial resistance (AMR), multidrug resistance (MDR) and virulence factors; siderophores (yersiniabactin and aerobactin), the hypermucoidy locus RmpADC, and combinations of aerobactin and RmpADC, which are indicators of a possible hypervirulent phenotype. The number of genomes per niche is displayed on/in the bars. **C) and D)** Proportion and ranges of heavy-metal resistance operons (HMRGs), plasmid replicon markers and thermoresistance genes. Statistical comparisons were performed using Kruskal-Wallis (overall) and Mann-Whitney (pairwise) tests for range, and chi-squared tests for proportions (overall and pairwise). Significance is denoted as follows: * P<0.05, ** P<0.01, *** P<0.001, ns P≥0.05.

**Fig. S3.**
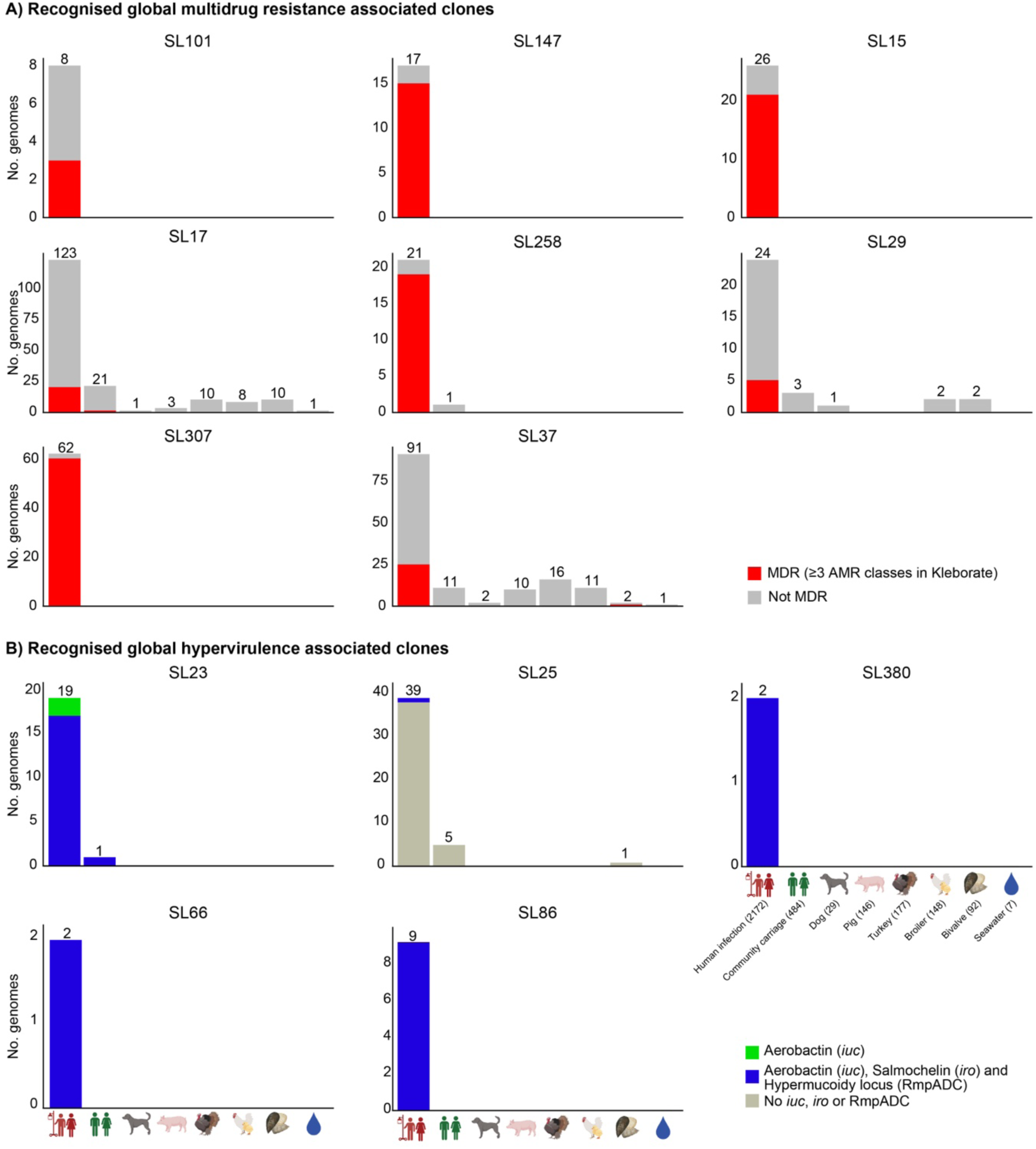
Recognised multidrug resistance (MDR) and hypervirulence associated clones. **A)** Number of genomes belonging to MDR associated clones (facet), shown by source and coloured by whether they were MDR (red) or not. **B)** Number of genomes belonging to hypervirulence associated clones (facet), shown by source and coloured by virulence loci that were present (inset legend). The total number of genomes per source with these features is indicated on top of the bars.

**Fig. S4.**
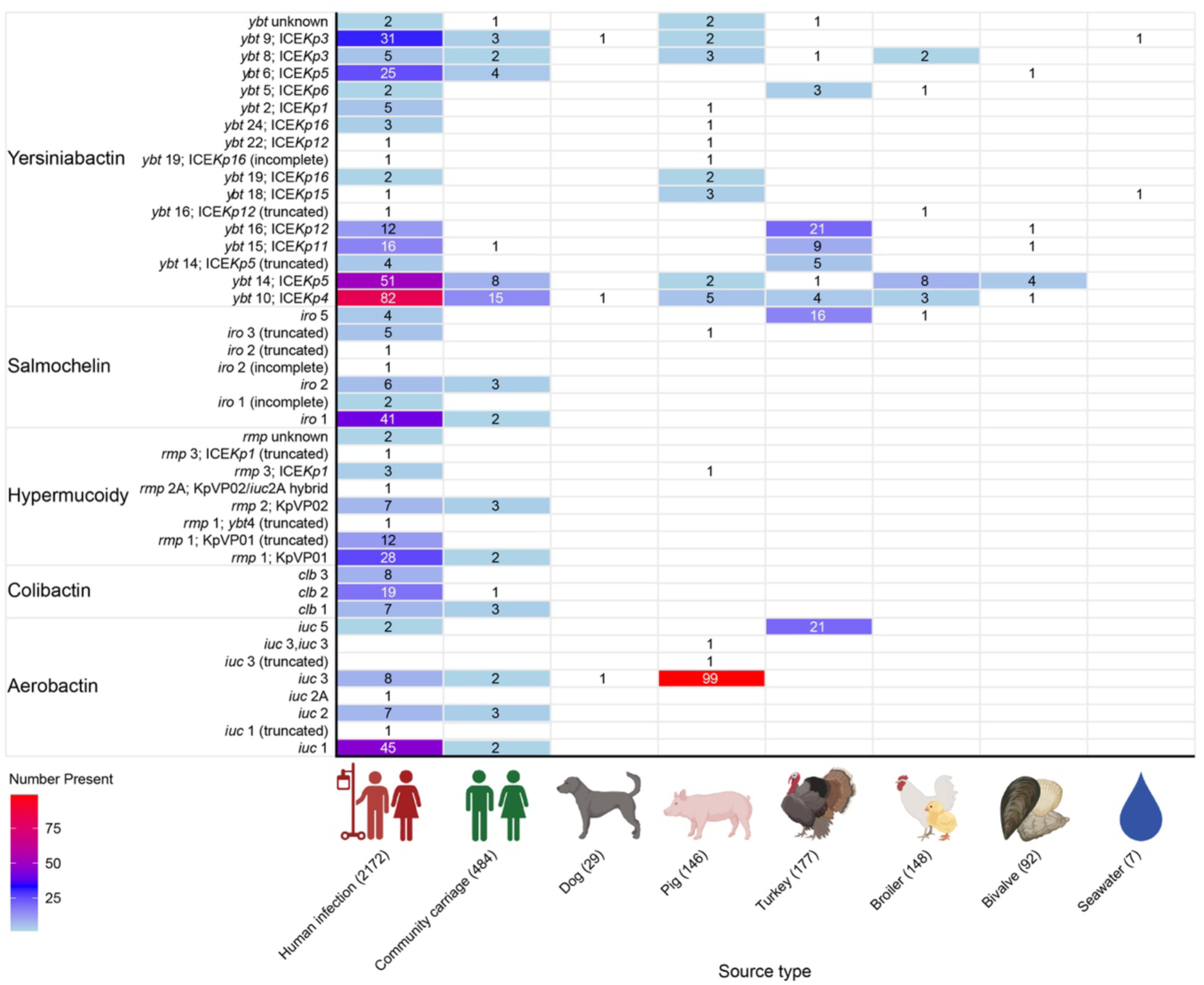
Virulence factors by source. The number of genomes per source (columns) that have the virulence locus specified in the rows. All loci are shown except for yersiniabactin, where only loci present in ≥1 non-human source are displayed (i.e. 17/63 loci).

**Fig. S5.**
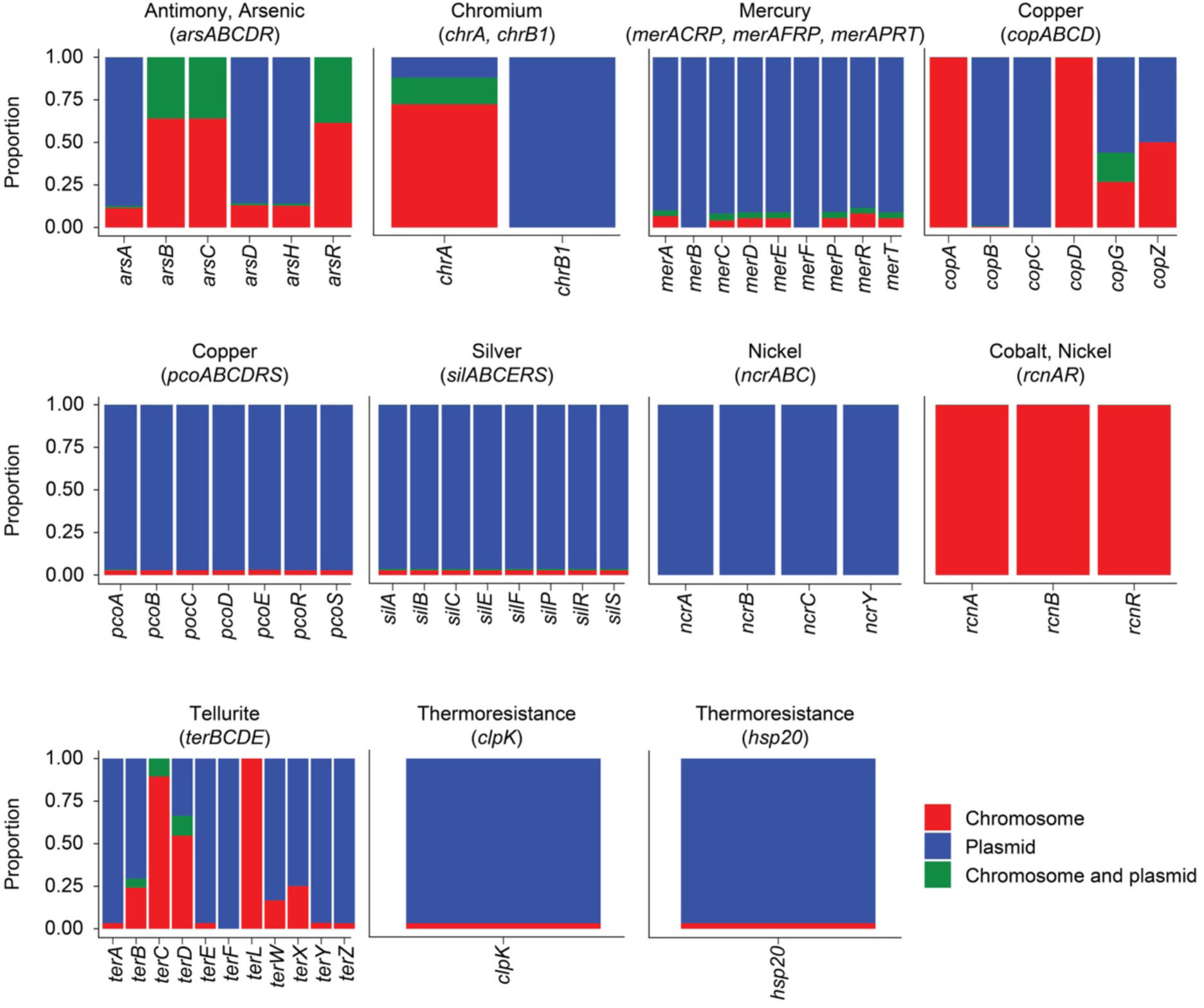
Distribution of heavy metal- and thermoresistance genes by replicon type. The closed genome collection of 550/3,255 genomes was utilised to determine if features were more commonly present on chromosomes or plasmid sequences. The proportions of genes are coloured by whether the gene was found on the chromosome (red), on plasmids (blue) or on both within the same genome (green).

**Fig. S6.**
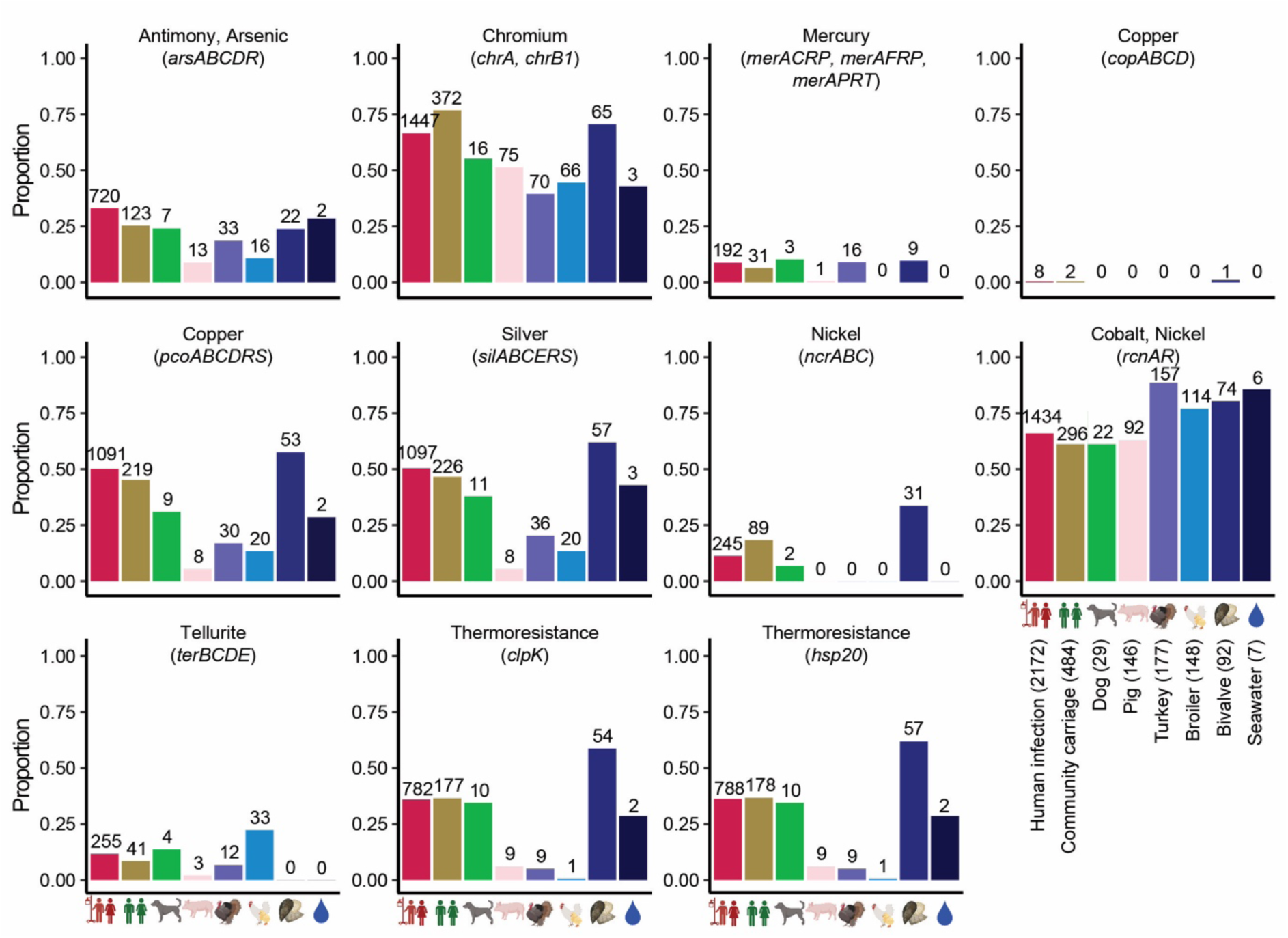
Distribution of heavy metal and thermoresistance genes by source. The facets show the presence of different heavy metal or thermoresistance genes or operons by source. The bars show the proportion of genomes by source; the number of genomes is indicated on top of each bar.

**Fig. S7.**
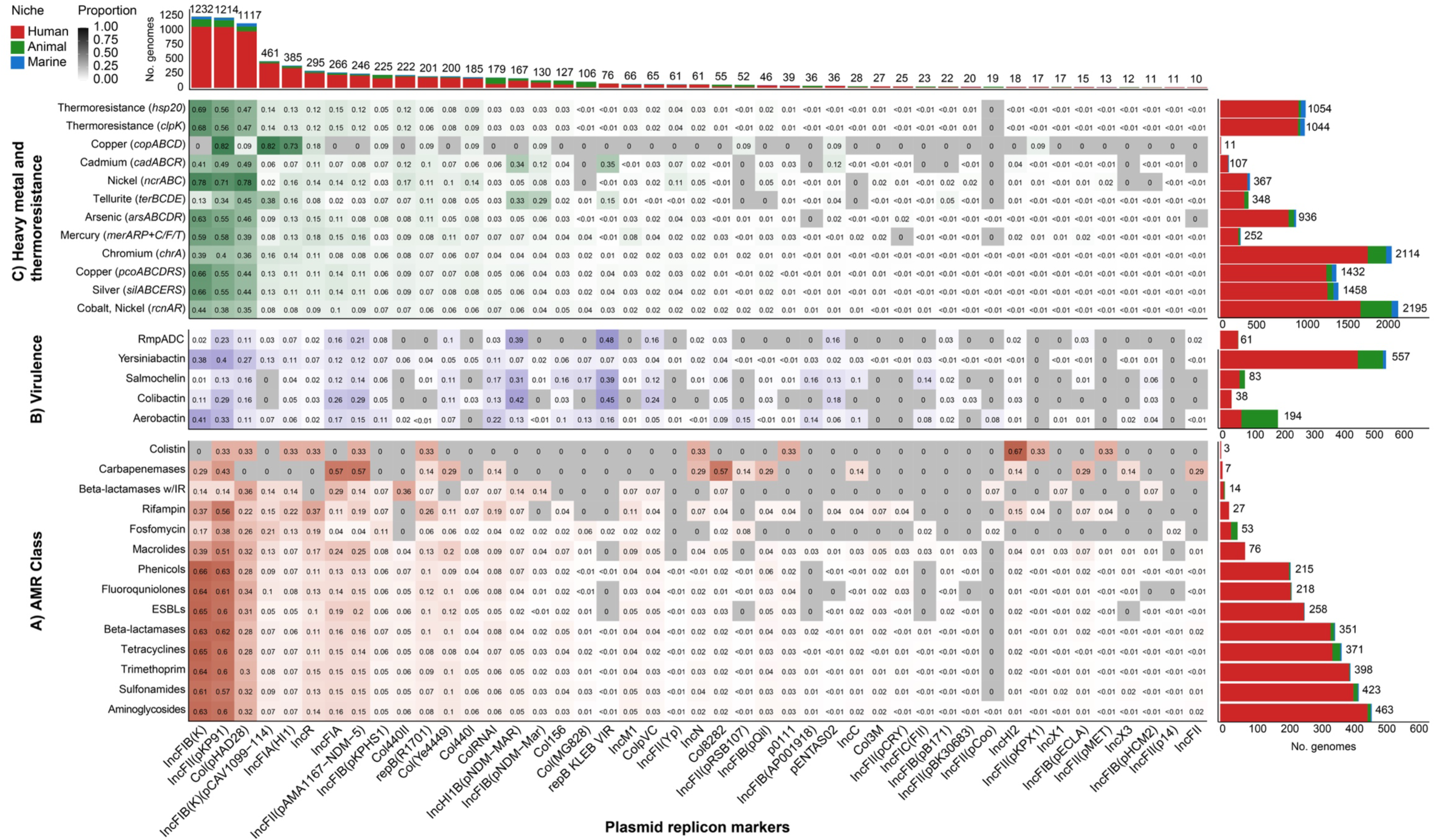
Plasmid replicon markers associated with clinically relevant features. Proportion of genomes with plasmid replicon markers (x-axis, showing markers found in ≥10 genomes each, i.e. 45 of in total 85 plasmid replicon markers) found together with **A)** Acquired antimicrobial resistance (AMR) determinants by class, **B)** Acquired virulence factors, and **C)** Heavy metal- and thermoresistance operons/genes. The marginal histograms show the number of genomes containing each feature, coloured by niche.

**Fig. S8.**
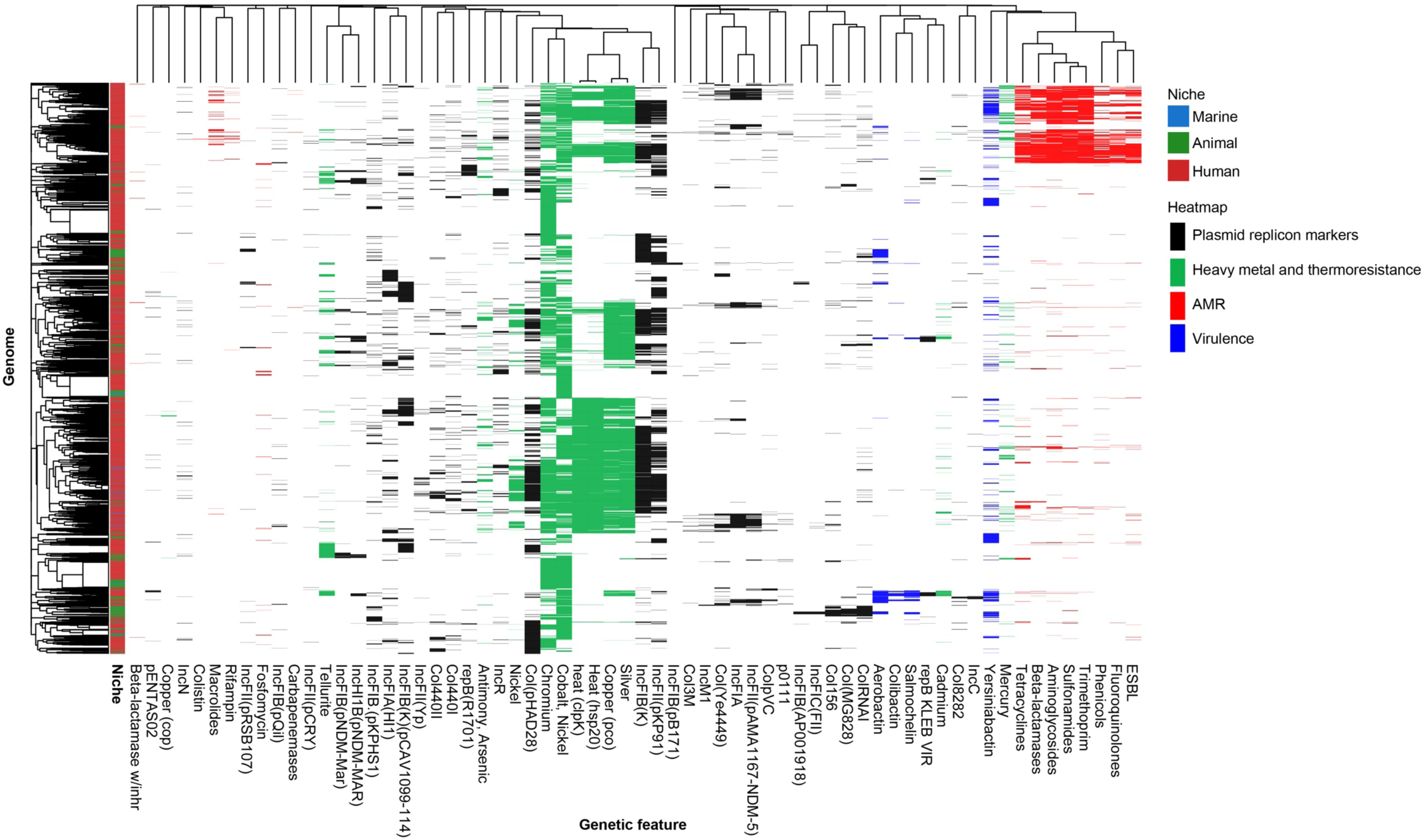
Co-occurrence of genetic features within genomes. A clustered heatmap showing the presence of genetic features (x-axis) in genomes (y-axis) and the niche they belong to (left-most column, as per inset legend). The features are grouped and coloured by: antimicrobial resistance (AMR) classes, virulence factors, heavy metal- and thermoresistance operons/genes, and plasmid replicon markers that were present in >20 genomes.

**Fig. S9.**
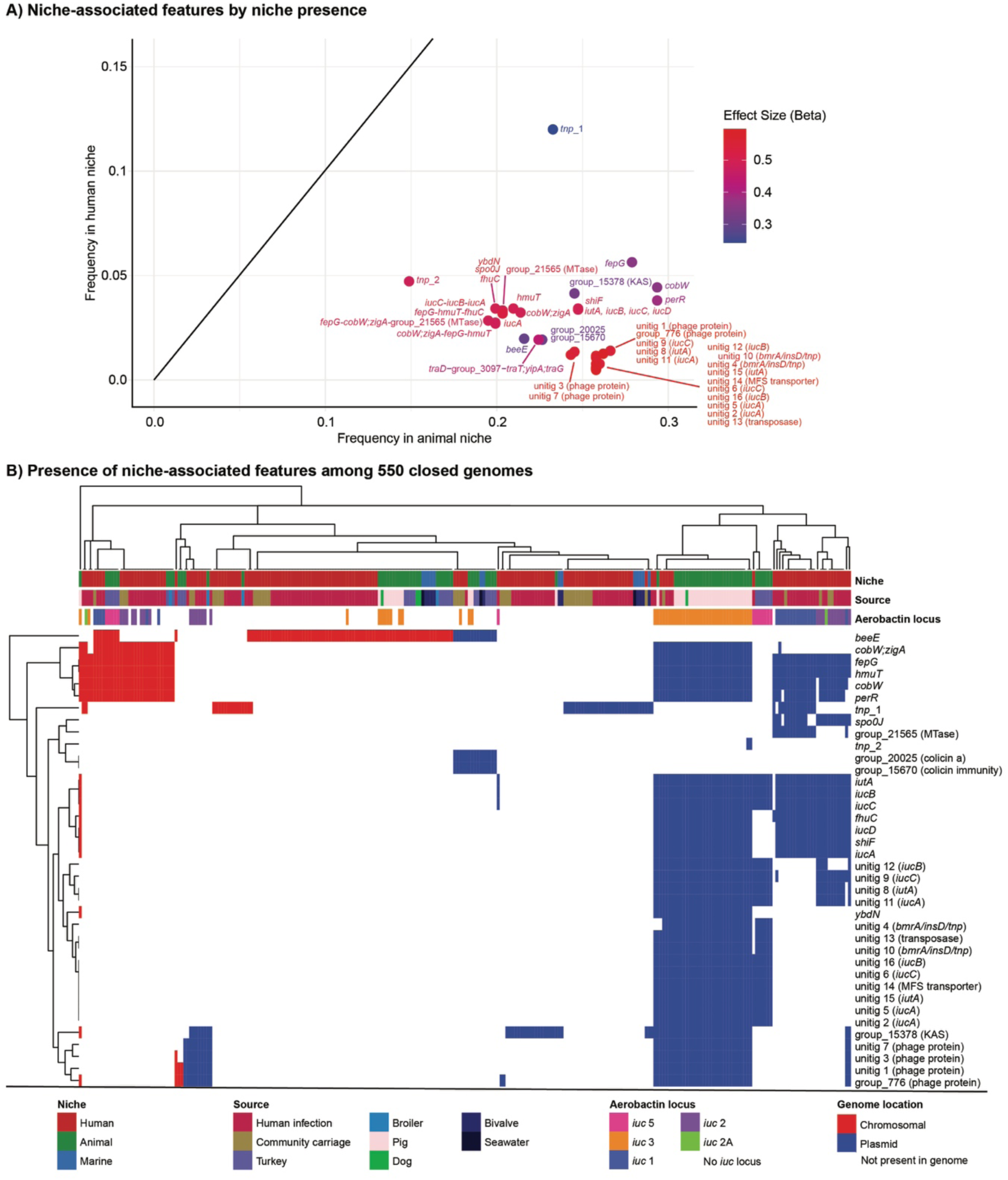
Presence of niche-associated genetic features by niche and replicon type. **A)** Niche-associated genetic features by their proportional presence in isolates with the phenotype (x-axis, i.e. in *K. pneumoniae* isolates from animals) or without (y-axis). **B)** Presence of significant features in the closed genome collection (n=550/3,255). The heatmap shows if the hits were found on chromosomes (red) or plasmids (blue). The three columns above the heatmap show niche, source and the presence of aerobactin (*iuc*) locus per genome (inset legend).

**Fig. S10.**
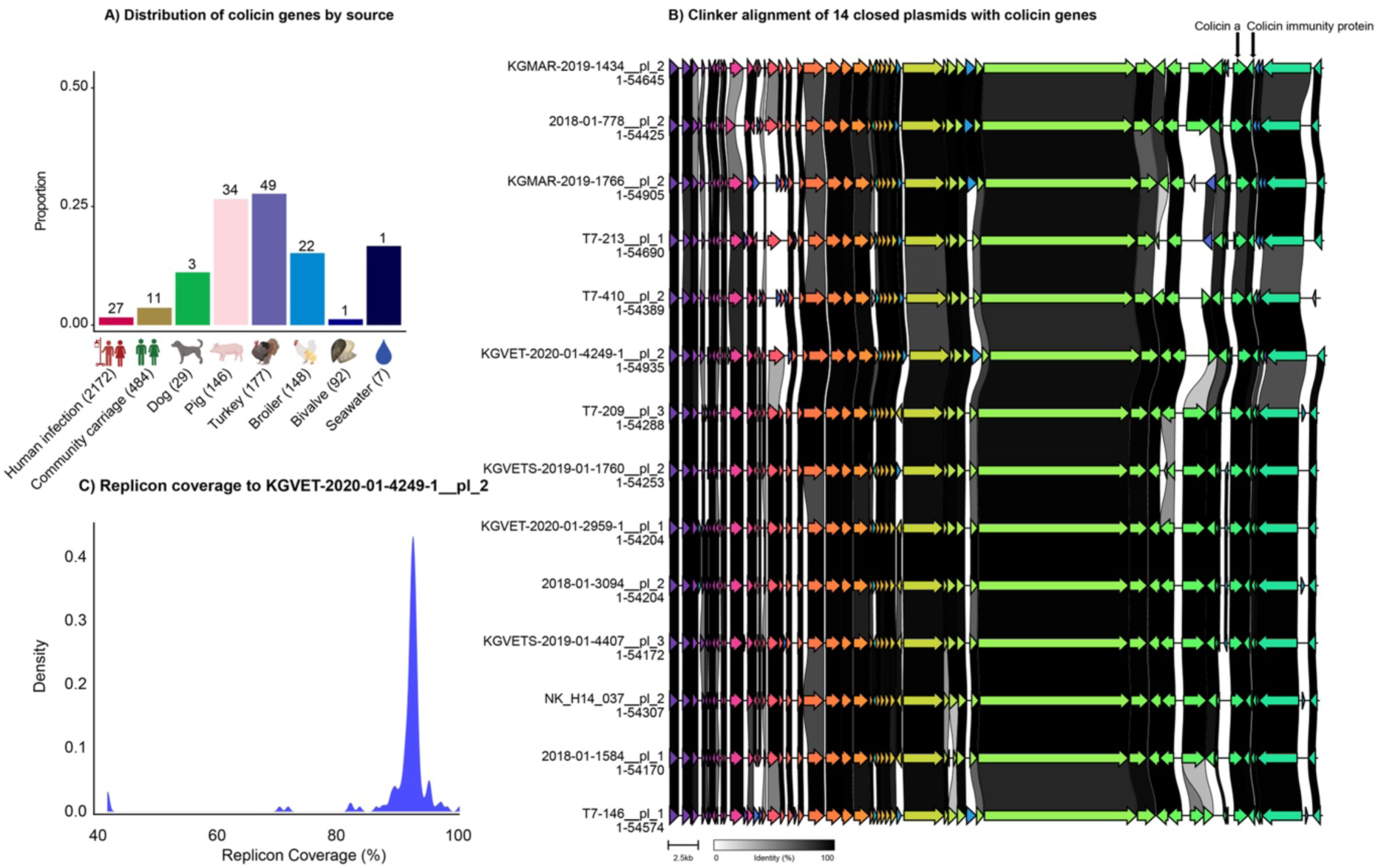
Distribution of colicin genes. **A)** By source; colicin a and colicin immunity protein were co-distributed in 148 genomes across the eight sources. **B)** Clinker-alignment of closed replicons encoding colicin. Fourteen of the 148 genomes had closed genomes and were aligned, revealing that the colicin genes (indicated on top of plot) were located next to each other on highly similar plasmids. **C)** Genomes with the colicin-encoding plasmid. The largest plasmid from B) was used as a reference to align the 148 short-read sequenced genomes against, revealing that the colicin-genes were found on highly similar plasmids in most genomes.

**Fig. S11.**
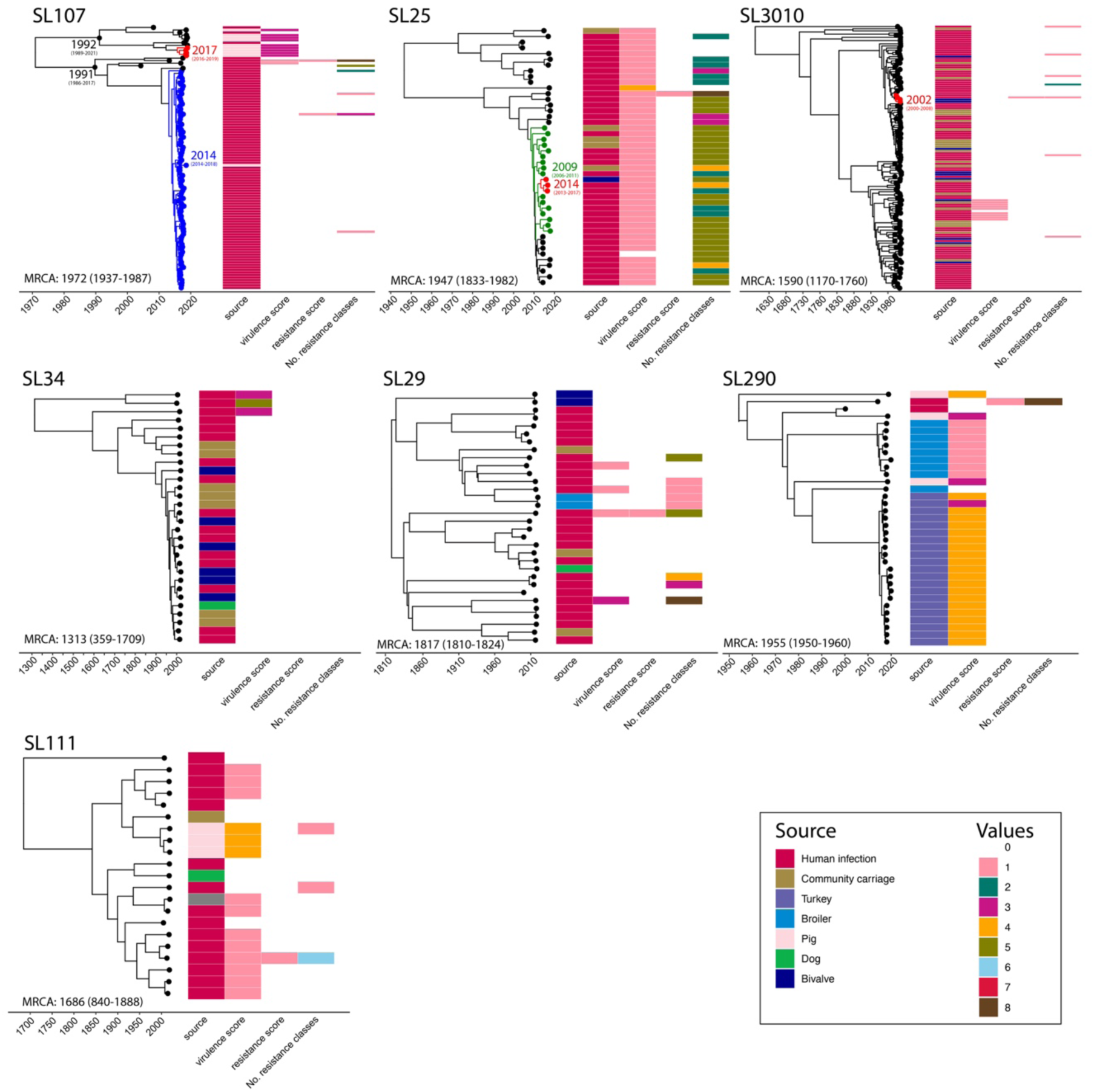
Dated trees of prevalent niche-overlapping SLs. The most prevalent niche-overlapping SLs were old with variable AMR and virulence content. The dated phylogenies are shown together with source information, virulence score, resistance score and the number of antimicrobial resistance classes. The most recent common ancestor (MRCA) of each SL is indicated with 95% HPD intervals. SL107, SL25 and SL3010 included cross-niche strain-sharing (shared ≤22 single nucleotide polymorphisms). The MRCAs of the genome pairs involved in those are indicated on the tips. The remaining dated SLs were frequent across the niches but did not include cross-niche strain-sharing at ≤22 SNP (see also Table S4).

**Fig. S12.**
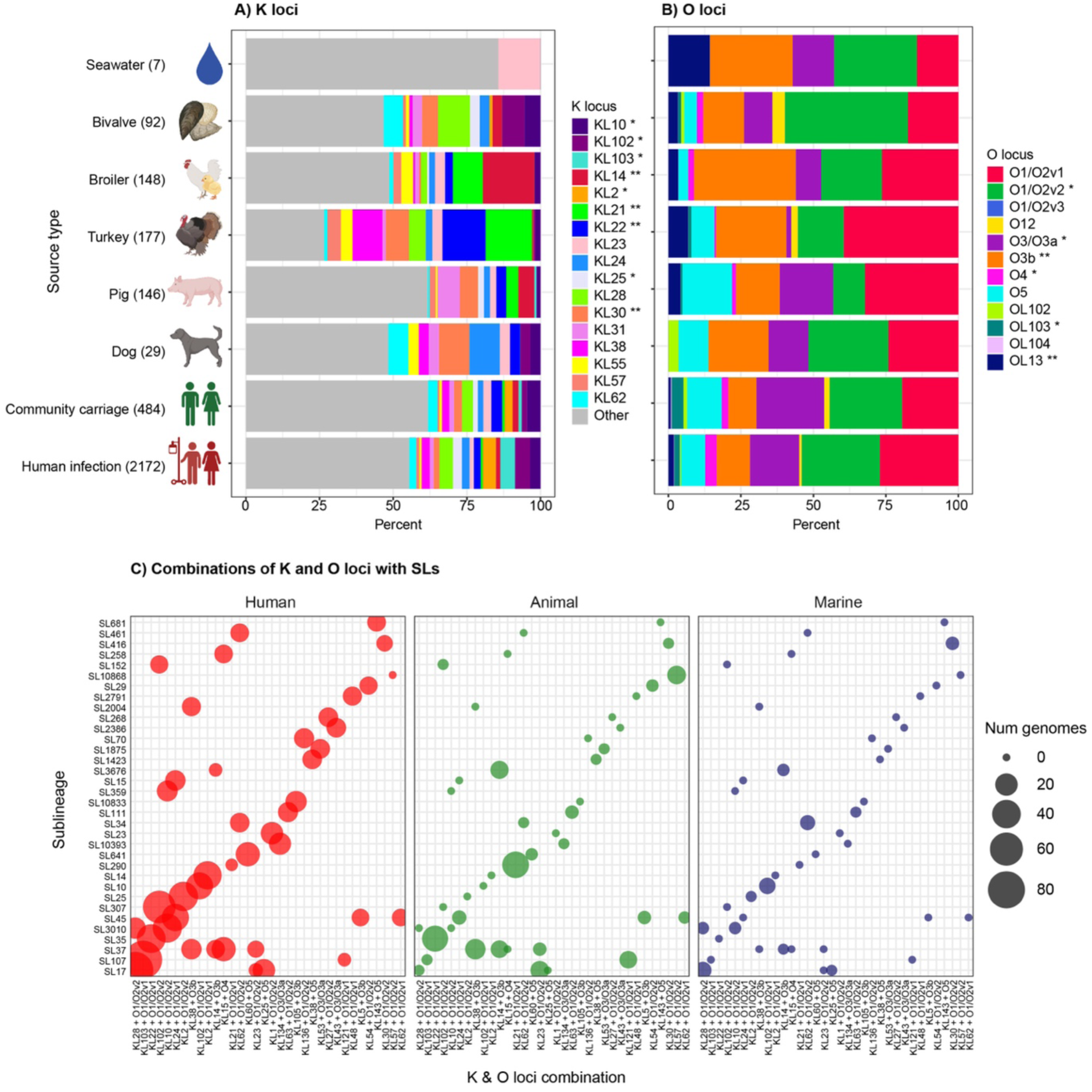
Capsule (K) and O loci by source and sublineages (SLs). **A)** Distribution of K loci among the sources. The 10 most prevalent K loci from each of the human and non-human samples are shown. **B)** Distribution of O loci among the sources. * K and O loci that were overrepresented among the human samples; ** loci that were overrepresented in the non-human samples (see Table S6). **C)** The most common combinations (>10 genomes) of K and O loci (x-axis) and SLs (y-axis). The bubbles indicate the number of genomes, and are shown for each of the three niches: human, animal and marine.

**Fig. S13.**
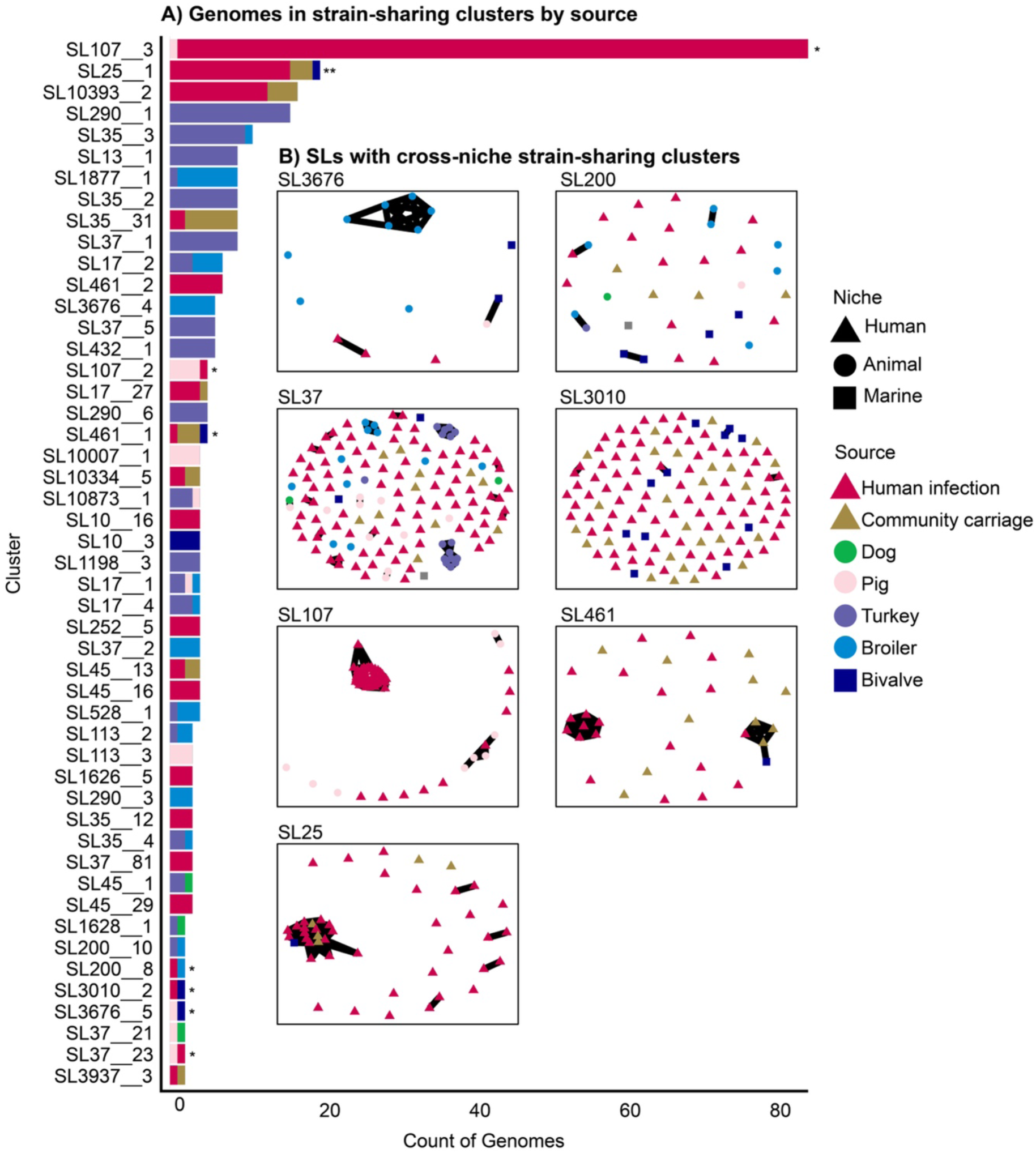
Strain-sharing clusters by source. **A)** Number of genomes within strain-sharing clusters (sharing ≤22 SNPs), coloured by source. All strain-sharing pairs are shown, except for clusters with only 2 genomes where they came from the same source (62 human infection, 9 community carriage, 8 pig, 4 turkey, 9 broiler and 13 bivalve pairs). To distinguish clusters within the same SL, cluster names were assigned using the SL and a sequential number. **B)** Seven SLs had ≥1 strain-sharing cluster between ecological niches (indicated with * in A). The nodes represent the genomes within each SL. Lines were drawn between pairs of genomes if they shared ≤22 SNPs. The nodes are shaped by niche and coloured by source (inset legend).

**Fig. S14.**
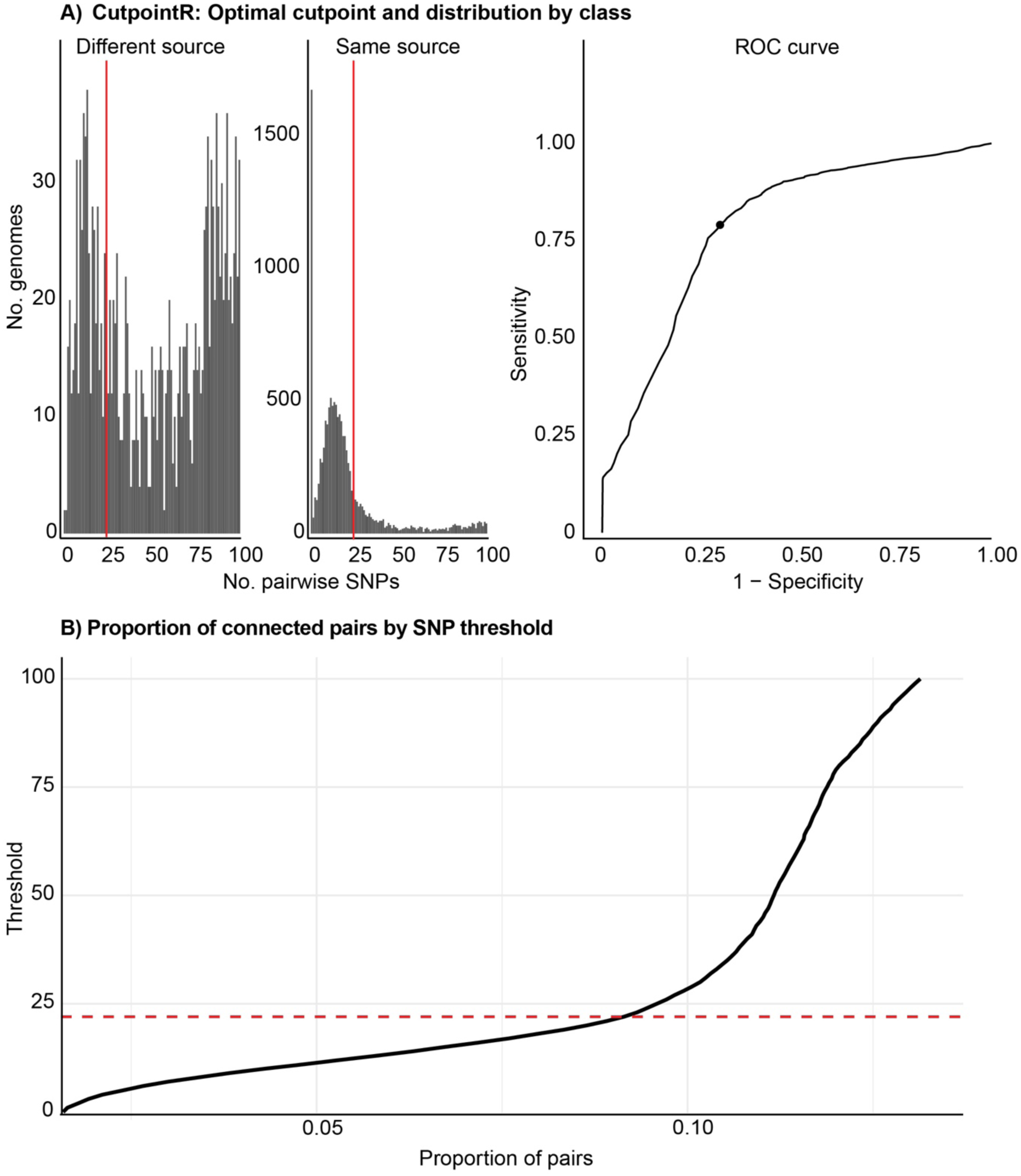
Strain-sharing pairs by single nucleotide polymorphism (SNP) thresholds. **A)** CutpointR was used to estimate an optimal SNP threshold of 22 (red line) for differentiating genome pairs within or between sources. **B)** The proportion of genome pairs (black line) given a SNP threshold (up to 100 SNPs). The red dotted line shows that the estimated optimal threshold of 22 is at an inflection point in the curve, suggesting it is an appropriate threshold for maximising the differentiation of strain-sharing pairs within or between sources.

**Fig. S15.**
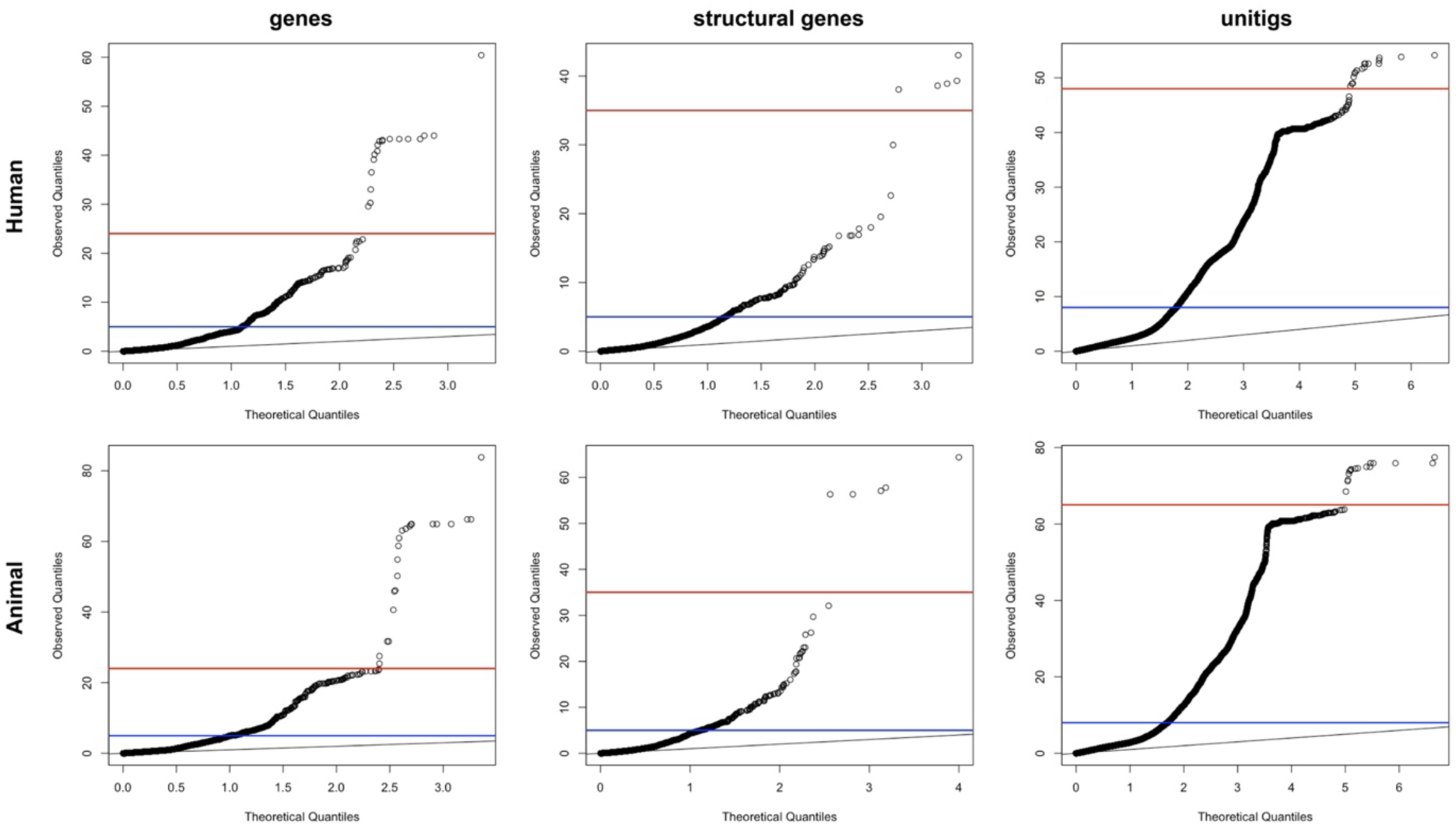
QQ-plots of the pyseer results. The blue line represents the p-value threshold suggested by pyseer, which was too high because the population structure had not been completely corrected for. The red line indicates a manually set p-value threshold, which was used instead, based on visual inspection of the data.

