## Supplementary Methods for "A genome-wide One Health study of *Klebsiella pneumoniae* in Norway reveals overlapping populations but few recent transmission events across reservoirs"

**Affiliations:**
[1] Department of Medical Microbiology, Stavanger University Hospital, Stavanger, Norway
[2] Department of Biological Sciences, Faculty of Mathematics and Natural Sciences, University of Bergen, Bergen, Norway
[3] Department of Medical Biology, Faculty of Health Sciences, UiT The Arctic University of Norway, Tromsø, Norway
[4] Research Section Food Safety and Animal Health, Department of Animal Health and Food Safety, Norwegian Veterinary Institute, Ås, Norway
[5] Institute of Marine Research, Bergen, Norway
[6] Biodiversity and Epidemiology of Bacterial Pathogens Unit, Institut Pasteur, Université Paris Cité, Paris, France
[7] Department of Infection Biology, Faculty of Infectious and Tropical Diseases, London School of Hygiene & Tropical Medicine, London, United Kingdom
[8] Department of Clinical Science, Faculty of Medicine, University of Bergen, Bergen, Norway
[9] Department of Infectious Diseases, School of Translational Medicine, Monash University, Melbourne, Australia
[10] Department of Microbiology and Infection Control, University Hospital of North Norway, Tromsø, Norway
[11] Norwegian National Advisory Unit on Detection of Antimicrobial Resistance, Department of Microbiology and Infection Control, University Hospital of North Norway, Tromsø, Norway
[12] Section for Bacteriology, Department for Analysis and Diagnostics, Norwegian Veterinary Institute, Ås, Norway
[13] Department of Microbiology and Immunology, University of Melbourne at the Peter Doherty Institute for Infection and Immunity, Melbourne, Australia

[b] These authors contributed equally

**Keywords**: One Health; *Klebsiella pneumoniae* species complex; genomics; ecology; zoonotic transmission

### Sample selection

We included in total 3,255 *Klebsiella pneumoniae* species complex (KpSC) isolates in this study. Isolates from human infections (n=1920) were collected from 22 hospital laboratories across Norway, between 2001 and 2018 [1,2]. Human faecal carriage isolates (n=484) were collected from a general adult population in the Tromsø municipality, Norway, during 2015-2016 [3]. From marine environments, *K. pneumoniae* isolates (n=99) were recovered from 7 surface seawater samples and 92 bivalves and sea urchins (in 2016, 2019 and 2020), but none were recovered from fish or sediment [4–6]. For bivalves and sea urchins, batches of 10-20 individuals were pooled to obtain an isolate. The samples from fish, surface seawater and sediment were from unique individuals or samples. Carriage (caecal or faecal) samples from pigs (n=146, collected in 2019), turkey flocks (n=113 in 2018, n=60 in 2020), broiler flocks (n=90 in 2018 and n=55 in 2020), wild boars (n=27 in 2020), dogs (n=16 in 2019) and cattle (n=12 in 2019) were collected via the Norwegian monitoring program for antimicrobial resistance (AMR) in the veterinary sector (NORM-VET) [7–9]. From turkeys and broilers, ten caecal samples were pooled from each flock to obtain one sample for screening. From pigs, dogs, cattle and wild boar only one sample per herd/animal was included. Additionally, 20 clinical isolates were included from individual dogs, turkeys and broilers. The dataset was divided into eight source types for comparison throughout the study: KpSC from human infection, community carriage, dogs, pigs, turkeys, broilers, marine bivalves and seawater.

### Identification of heavy metal operons and thermoresistance genes

We searched the annotated assemblies for experimentally confirmed heavy metal resistance genes listed in the Antibacterial Biocide and Metal Resistance Genes (BacMet) database (http://bacmet.biomedicine.gu.se/index.html) [10]. Heavy metal resistance is typically encoded by operons of genes, where certain genes must be present for the operon to be active. This has been experimentally confirmed for some metals in some bacterial species, but far from all. Based on literature searches, we used the following operons to determine presence or absence of heavy metal resistance: Arsenic resistance was determined by the presence of at least *arsABCDR* [11-13]. Resistance to chromium was defined by the presence of *chrA*. The plasmid-encoded *chrB1* is believed to increase resistance but is not essential [11,14]. Mercury resistance was determined by the presence of at least *merAPR* and either *merC, merF, or merT* [11,15]. Copper resistance operons were defined as *cusABCF*, *pcoABCDRS* or *copABCD* [11,16]. Resistance to nickel was defined as the presence of *ncrABC* [17]. Resistance to cadmium was defined as *cadABC* [13,18,19]. Silver resistance was defined as the presence of at minimum *silABCERS* [11]. Several operons were used to determine tellurite resistance, including *klaABC*, *kilA* and *telAB*, *tehAB,* and *terBCDE* [20]*.* The presence of *zitB* genes indicated resistance to zinc, and the *zntAR* operon defined resistance to cadmium, lead, and zinc [21,22]. The *czcCBA* operon was used to detect the multi-metal efflux pump for cadmium, zinc, and cobalt [11,18]. The presence of *rcnAR* defined resistance to cobalt and nickel [23]. There were no complete operons of *klaABC, kilA, telAB, czcABC,* or *cusABCF* in any genomes in the overall collection and they were therefore not shown in figures/data. The following operons were present in ≥99% of genomes in all niches and were therefore excluded from the figures: *cadABC* (n=3244), *cueOR* (n=3249), *tehAB* (n=3254), *zitB* (n=3253), and *zntAR* (n=3245). Thermoresistance was defined as the presence of the genes *clpK* or *hsp20* [24,25]. We utilised our hybrid genome assembly collection (n=550/3,255) to determine if these genes/loci were most commonly encoded on plasmids or on chromosomes (Fig. S5).

### Inferring cross-talk between niches

Of the 107 niche-overlapping SLs, we selected those that were represented by ≥20 genomes and had been collected over at least a 5-year period (n=15 SLs). We first estimated dated phylogenies using both our local dataset and publicly available genomes for robust clock rate estimates. We then estimated dated phylogenies of the local datasets only, using the best fit clock models from the global phylogenies, and setting the mutation rates identified from the global phylogenies as the initial rate of substitutions per genome.

For each SL, we downloaded all publicly available genomes (short-reads) with known year of collection identified on https://pathogen.watch on 01.07.2023. They were assembled with the same methods as the local genomes (TrimGalore v0.6.7 [https://github.com/FelixKrueger/TrimGalore] and assembled with SPAdes v3.15.4 [26]). This gave a total number of 2,974 public and 1,020 local genomes for these analyses. For each SLs, both the public and local genomes were included to identify and filter recombinations using verticall v0.4.2 (https://github.com/rrwick/Verticall). The output was used by BactDate v1.1.1 [27] to infer dated phylogenies of each SL. We specified exact dates of sampling for genomes where this was available (n=707 from our local dataset), and for genomes where only sampling year was available we specified a year range (n=313 from our dataset, all of the public genomes) (Table S5 lists the accessions and relevant metadata).

BactDating was run with three clock models, a strict ("strictgamma"), mixed ("mixedgamma") and relaxed log normal clock model ("relaxedgamma"), with a constant coalescent demographic model. Each model was run in three independent replicates, to $10^8$ Markov Chain Monte Carlo (MCMC) iterations, until all parameters reached >200 effective sampling size (ESS). For each SL, the best fit model was determined using the modelcompare function in BactDate.

Prior to running BactDate, we tested for temporal signal (association between time and genetic divergence) using the root-to-tip regression function in BactDate with the verticall tree. The root-to-tip genetic distances were positively associated with the dates of isolation (Table S4), indicating molecular clock signals. After running BactDate, the statistical significance of the temporal signal was tested by running the same model again but with sampling dates set equal, and then comparing the two models.

To assess the relatedness of respectively colicin-containing and *iuc*3-containing plasmids across the niches, RedDog v1beta.11 (https://github.com/katholt/RedDog) was used to align all 3,255 short-read genomes against the largest closed colicin-containing (KGVET-2020-01-4249-1, 54.9 kbp) and *iuc*3-containing (KGVETS-2019-01-1806, 148.4 kbp) plasmids from the collection and comparing the number of SNPs and replicon coverage. To compare the structure of the colicin-containing genomes, clinker v0.0.29 [28] was used to align all closed colicin-containing plasmid sequences.

### Assessing niche enrichment

To assess niche-enrichment, we performed genome-wide association studies (GWAS) with pyseer v1.3.11 [29]. We looked for associations using four genetic features: 1) the gene presence absence matrix from panaroo v1.3.3 [30], 2) the structural presence absence matrix from panaroo, 3) SNPs generated with RedDog mapping against the reference SGH10 (GenBank accession no CP025080.1, re-annotated here with Bakta v1.8 [31]), and 4) unitigs generated with unitig-caller v1.3.0 (https://github.com/bacpop/unitig-caller). The analysis was restricted to *K. pneumoniae* due to differences in KpSC species distribution and counts in the three niches.

We used the linear mixed model (LMM) in pyseer, accounting for population structure by using a maximum likelihood tree generated by IQ-tree v2.2.6 [32] from the same alignment as we derived the SNPs from above. The pyseer-script 'phylogeny_distance.py' was used to midpoint root the tree and to calculate the distance matrix needed for population structure correction. For pyseer, we set the minimum allele frequency to 5% and the maximum to 95%, to exclude very rare variants and variants that were very common in the dataset from the analysis. QQ-plots of the pyseer results revealed that population structure had not been sufficiently accounted for. We therefore manually decreased the p-value thresholds for significant associations based on the QQ-plots (Fig. S15). Genes, structural genes and SNPs were identified based on the Bakta-annotations. Unitig locations were identified by searching the Bakta-annotated hybrid genome assembly collection (n=550/3,255).
